# Modelling of tuna around fish aggregating devices: the importance of ocean flow and prey

**DOI:** 10.1101/2022.09.20.508652

**Authors:** Peter D. Nooteboom, Joe Scutt Phillips, Christian Kehl, Simon Nicol, Erik van Sebille

## Abstract

Catch and distribution of tuna in the ocean are typically investigated with ocean basin-scale models. Due to their large scale, such models must greatly simplify tuna behaviour occurring at a scale below ∼100 km, despite interactions at this level potentially being important to both catch and distribution of tuna. For example, the associative behaviour of tuna with man-made floating objects, that are deployed by fishers to improve their catch rates (Fish Aggregating Devices; FADs), are usually ignored or simplified. Here we present a model that can be used to investigate the influence of tuna dynamics below the ∼100 km scale on larger scales. It is an Agent-Based Model (ABM) of a hypothetical, tuna-like species, that includes their interactions with each other, free-floating FADs and prey. In this ABM, both tuna and FADs are represented by Lagrangian particles that are advected by an ocean flow field, with tuna also exhibiting active swimming based on internal states such as stomach fullness. We apply the ABM in multiple configurations of idealised flow and prey fields, alongside differing interaction strengths between agents. When tuna swimming behaviour is influenced equally by prey and FADs, we find that the model simulations compare well with observations at the ≲ 100 km scale. For instance, compared to observations, tuna particles have a similar stomach fullness when associated or non-associated to a FAD, tuna colonize at similar timescales at FADs after their deployment and tuna particles exhibit similar variations in continuous residence times. However, we find large differences in emergent dynamics such as residence and catch among different flow configurations, because the flow determines the time scale at which tuna encounter FADs. These findings are discussed in the context of directing future research, and an improved interpretation of tuna catch and other data for the sustainable management of these economically important species.

## 1. Introduction

Tropical tuna species provide some of the largest catches of high-trophic fish in the world, and as such the assessment and management of the fisheries they support are critical to ensuring sustainable stocks, food security, and livelihoods [1, 2] (FAO SOFIA). In the case of tropical tunas, the use of drifting Fish Aggregating Devices (FADs) by industrial purse seiner fisheries has markedly changed the efficiency within fishing grounds [3, 4]. FADs are floating objects (i.e. drogued buoys) that aggregate pelagic fish around them. Tens of thousands of FADs are deployed in the equatorial regions of the world’s oceans annually [5]. They usually include a GPS tracking system and generally an echo sounder to measure the biomass of surrounding fish. This increases the knowledge of where fish are most abundant. Therefore, FADs act as a peculiar trait of a predator by attracting tuna and other fish, where-after tuna are caught by purse seiners on or near the FADs. While it remains unclear whether tuna species are directly or indirectly attracted towards FADs [6], it is clear that they impact tuna behaviour and distribution [7, 8, 9, 10, 11].

To predict the distribution and abundance of tuna biomass in the oceans, models are often used, typically integrating catch and other fisheries data, and they are even coupled to ocean-biogeochemical models in some cases [12, 13, 14, 15, 16]. These population dynamics models use density-dependent functions as an abstraction of individual tuna behaviour, such as their foraging behaviour [12, 15], predator-prey equations, or trophic functions, that describe how the density of multiple species grows or declines in relation to one predating the other [17, 18, 19], or density-dependent catchability, describing the relationship between the abundance of a fish population and how easy it is to catch them for a given effort [20].

Although FADs have a substantial impact on tuna behaviour, distribution and catch [8], most tuna distribution models do not consider the direct interaction of tuna with FADs [12, 13, 14, 15, 16]. In contrast, they abstract their behavioural effect through separated fisheries with differing units of effort, and assumed catchability and selectivity parameters. Moreover, a lack of observations and models exists to test the assumptions that tuna models typically use at their sub-grid scale (i.e. ≲ 100 km), at which FADs introduce relevant dynamics for the distribution of tuna.

Observations of individual tuna [21, 22, 23, 24] and aggregated biomass [25, 26, 27] show variable patterns of colonisation and residence around FADs, and some of these dynamics have been replicated in simulation experiments of individual-based models [28, 9], which can have included the interaction between fish and FADs [9]. In contrast to Eulerian models [12, 13, 14, 15, 16], individual-based models allow for an individual-based quantification of tuna behaviour that can be compared to individual-based observed data [29]. Although often computationally more expensive compared to Eulerian models, individual-based models provide a ‘bottom-up’ approach, which implements the behaviour of individuals below the ∼100 km scale to obtain a better understanding on the emergence of complex predator-prey dynamics at large spatial scales [30, 31]. Furthermore, trophic functions have been shown to emerge from individual predator-prey dynamics [32, 33], and be responsible for complex and chaotic behaviour when involving multiple groups [34].

Recent development of particle-particle and particle-field interaction functionalities in the Parcels Lagrangian framework [35] allows us to extend the approach of previous tropical tuna individual-based models [28, 36, 37], to include the interaction of tuna with prey, FADs and ocean flow. Here we present a specific type of individual-based model (which we refer to as Agent Based Model; ABM), where the agent represent a group of tuna individuals [38, 39]. The ABM considers those dynamics that impact the distribution of tuna below the ≲ 100 km scale, and we apply the ABM in idealised configurations. The ABM enables us to test whether these interactions are relevant to explain specific observations that occur at this scale, such as the colonization and residence times at FADs, and tuna stomach fullness, and the extent to which their observed variability can be caused by ocean flow, prey dynamics and fishing strategies. This information could be used to improve population dynamics models and the decisions that managers base on their simulations. We explore potential mechanisms that lead to the simulated emergence of these dynamics.

## 2. Methods

### 2.1. Biological Assumptions

The temporal and spatial scales of observed tuna-FAD interactions are typically days to weeks and sub ∼ 1° × 1°, respectively [24, 40, 41]. It is therefore not necessary to include all dynamics that are relevant for the distribution of tuna in the ocean-basin scale, but rather to capture the local-scale interactions that may be responsible for the patterns observed. To create a minimally-appropriate behavioural model for a tuna-like species around floating objects, we have drawn on *in-situ* observations and assumed that tuna are motivated by feeding and avoiding predators [42].

Tropical tuna, at the size-classes that typically associate with FADs, are a schooling and aggregating species [8]. Here we consider only those species and size classes of tuna that interact with FADs in the surface, epipelagic layer, and hence do not include their vertical behaviour. Schooling provides fitness benefits from increased foraging success, genetic diversity and protection from predators [43]. Rather than include schooling dynamics directly, here we include attraction between tuna particles as a mechanism by which aggregations of schools may form in the absence of other drivers.

Tuna foraging behaviour and its dependence on the temporal evolution of their stomach fullness are complex [9]. However, alongside survival and reproduction, feeding is a fundamental driver of animal movement [44], potentially impacting tuna-FAD dynamics at the school level [8], and so simple foraging and hunger-driven mechanisms are included in our model. For foraging, we assume that tuna are attracted by the presence of their prey, following local gradients of prey density [29, 45]. Prey are consumed by tuna, allowing density-dependent feedback mechanisms. Every tuna particle has a stomach fullness, which represents the average stomach fullness of the tuna organisms that a particle represents. We assume that tuna particles forage when their stomach fullness is less than approximately 40%. Their stomach fills with a linear rate of 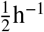 if food is available and their linear gastric evacuation rate is 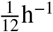, which broadly matches controlled experiments on tropical tuna species [46, 47].

The mechanisms behind tuna association near FADs have so far remained unclear [8]. However, empirical indications exist that FADs attract tuna particles when their distance is below ∼ 10km [11, 7]. Therefore, we base the attraction of our tuna to FADs on this value.

Apart from prey, the model presented in this paper does not include the effect of abiotic habitat on the tuna behaviour. This implies that we assume that habitat (such as temperature) is suitable for the tuna to survive throughout the studied domain. Moreover, we assume that tuna birth and mortality do not play a relevant role at the simulation timescale (i.e. 100 days) applied here. The size of tuna may have an influence at these scales (i.e. 100 days), since different sizes of tuna may have a different attraction strength towards FADs. Although FAD attraction strength is a parameter in the model, for simplicity, we assume that there are no ontogenetic changes to tuna behaviour during the course of a simulation.

### 2.2. Model

In the two-dimensional model, we use a rectangular domain Ω = [0, *L*_*x*_] × [0, *L*_*y*_]. We consider two types of particles (representing tuna and FADs) and one type of field that represents prey. The tuna particles interact with the prey field, the FADs and with each other. The prey field passively interacts with tuna particles through depletion. FAD particles interact with tuna particles through attraction and depletion.

In order to reduce computational costs of simulations, a tuna particle is not an individual tuna fish, but rather an entity representing a minimum group of tuna [29]. Since these particles represent a minimum group of tuna and not individual organisms in this paper, we refer to this model as an Agent-Based Model (ABM) instead of an ‘individual-based model’. Similarly, we abstract individual prey to a Eulerian field to minimize computational overhead. This field drives the active searching behaviour of individual tuna through random and directed movements, which have been shown to result in similar density evolution through time to diffusion- and taxis-like processes used to model the movement of animals [12, 48, 29, 45].

Each FAD particle *j* (*j* = 0, …, *F*) is passively advected by horizontal ocean currents. It is displaced every time step Δ*t*:

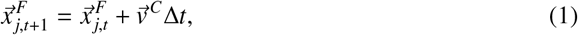

where 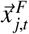 is the two-dimensional location of FAD *j* at time *t* (*t* = 1, …, *T*) and 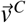 is the ocean flow velocity. Every tuna particle *i* (*i* = 1, …, *N*) is also advected by ocean currents, similarly to the FADs. However, tuna particles also swim. Overall, their trajectories are governed by:

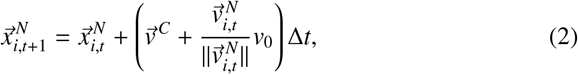

where 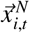 is the location of tuna particle *i* at time *t*. Similarly to [48], the magnitude of the swimming velocity is deterministic, given by 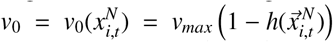, where *v*_*max*_ determines the maximum magnitude of the swimming velocity and *h*(*x*) is the prey index at location *x*, which is nearest-interpolated from the prey index field *h* : Ω × [0, *T*] → [0, 1], having a resolution of Δ*x*. As a result of this implementation, tuna particles swim faster if the prey abundance index *h* is lower.

The swimming direction of tuna particles is determined by:

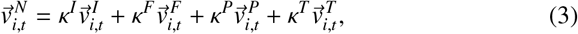

where 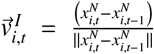 represents inertia [32]: tuna are more likely to keep swimming in the same direction compared to other directions. The parameter values *κ*^*I*^, *κ*^*F*^, *κ*^*P*^, *κ*^*T*^ ≥ 0 determine the relative contributions to the tuna swimming direction from inertia, FADs, prey and other tuna particles, respectively. Hence, the most dominant dynamics that determine the swimming direction of tuna can be easily tuned with these four parameters, of which the latter three are described below.

First, the tuna swimming direction towards FADs is given by:

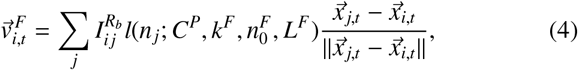

where *n*_*j*_ is the number of tuna that are closer than a distance *R*_*a*_ to FAD *j* (these tuna particles are ‘associated with’ FAD *j*). 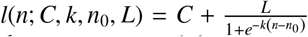 is the logistic function. Hence, the more tuna particles are associated with a FAD *j*, the stronger the swimming direction of tuna particles is determined by FAD *j* compared to other neighbouring FADs, creating a positive feedback for attraction to FADs [49]. The value of *R*_*b*_ is the interaction distances between tuna and FAD particles:

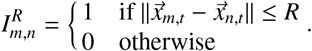

Second, the trajectories of tuna particles depend on the interactive prey index field *h*. The interaction between tuna particles and the prey field imply that tuna depletes the prey field every time step if prey is locally available, with the value 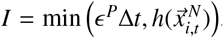. The total number of prey in the domain remains constant, since depleted prey is redistributed at another location in the domain. This location is determined by a probability density function, such that it is more likely that prey is added at a location where the prey index was large at *t* = 0. As a tuna particle *i* depletes the prey field, it reduces the stomach emptiness *St*_*i,t*_ ∈ [0, 1] at time *t* [9]:

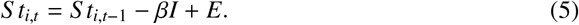

Here *E* = min (*ϵ*^*E*^ Δ*t*, 1 − *St*_*i,t*_) is the evacuation rate of the stomach.

Tuna swim towards high concentrations of the prey field through a taxis behaviour according to [48]

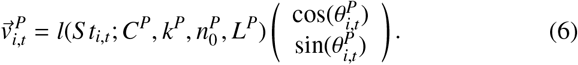

Here 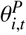 is drawn from the ‘von Mises’ distribution, with mean *θ*_0_ (which has the same direction as the gradient of the prey field, ∇*h*) and concentration parameter *κ*^*M*^ = *α* ∥∇*h*∥. Hence, the standard deviation of the von Mises distribution is lower if ∇*h* is higher. The logistic function *l*(*S t*_*i,t*_) in Eq.6 implies that the swimming direction of tuna is more strongly determined by the prey index gradient if their stomach is emptier [9].

Third, tuna particles are attracted towards each other if *κ*^*T*^ > 0:

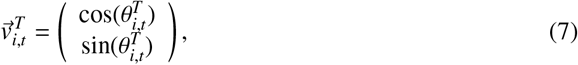

where 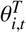 is drawn from the von Mises distribution with mean the direction of 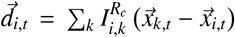 and concentration parameter 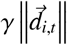, where *R*_*c*_ determines the interaction distance between tuna particles.

To summarize, stochasticity is included to the model in three ways. First, the level of stochasticity of the tuna swimming towards high prey abundance is controlled by *κ*^*m*^. Second, stochasticity of the tuna swimming direction towards other tuna is determined by *γ*. Third, the initial location of both FADs and tuna particles is random.

### 2.3. Fishing strategies

To examine the effect of differing model configurations on an idealised catch of tuna, fishing effort was kept constant at a single fishing event each day in the simulations. If a tuna particle is caught during such an event, it is removed and released at a random location in the domain, in order to keep the tuna density constant in the domain [32]. Four contrasting fishing strategies were implemented (Table 1). The first strategy (FS1) is based on fishing of FAD unassociated tuna. In this strategy, fishers have no information about any FAD, but they use sonar and sometimes helicopters to locate tuna [50]. Hence, we assume for our simulations that they simply search the domain to locate schools of tuna, performing a single fishing event near a random tuna particle each day (i.e. all tuna particles are caught with a probability *ϵ*^*T*^, if their distance with the randomly picked tuna particle is lower than *R*_*a*_).

**Table 1:**
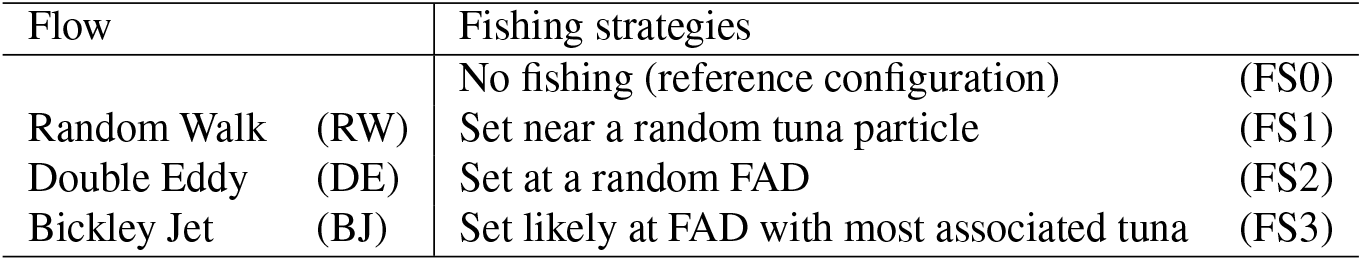
Overview of tested flow configurations and fishing strategies

| Flow |  | Fishing strategies |  |
| --- | --- | --- | --- |
|  |  | No fishing (reference configuration) | (FS0) |
| Random Walk | (RW) | Set near a random tuna particle | (FS1) |
| Double Eddy | (DE) | Set at a random FAD | (FS2) |
| Bickley Jet | (BJ) | Set likely at FAD with most associated tuna | (FS3) |

The second type of fishing strategy assumes that fishers know the location of all FADs. They choose a FAD to set their nets and catch every associated tuna particle with a probability *ϵ*^*T*^. A parameter *p* ∈ [0, 1] determines the extend of the fisher’s information on which FAD has most tuna associated with it. We order the FADs *j* = 0, …, *F* from high to low number of associated tuna. The probability that a fishing event occurs at FAD *j* is given by the geometric distribution:

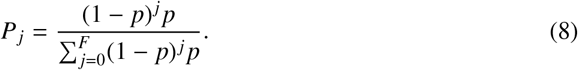

Hence, for *p* = 1, fishers have complete information on the number of associated tuna at all FADs, and always set at the FAD that has most associated tuna. For *p* = 0, the fishers have no information about the number of associated tuna at FADs, and choose a FAD with equal (i.e. uniform) probability. In this paper, we test fishing strategies *p* = 0 (FS2) and *p* = 0.95 (FS3).

We also use a reference fishing strategy (FS0). In FS0, no fishing events occur and no tuna particles are caught in the simulations.

### 2.4. Particle-particle interaction in Lagrangian simulations

Parcels is a framework for computing virtual Lagrangian particle trajectories in ocean flow [51]. For this study, we developed a novel ABM method with the Lagrangian particle advection for physical oceanography, which includes the interaction between these virtual particles (i.e. tuna and FADs).

Inspired by established procedures for *smoothed particle hydrodynamics (SPH)* for particle-based fluid-flow [52, 53], a specialised procedure of three-dimensional kD-Tree construction [35], as is common also for SPH simulations [54], was integrated into the Parcels framework. The focus of the developed method is on rapid rebuild- and particle indexing in a threedimensional geospatial coordinate frame. In global ocean simulations over long time spans, the arbitrary timestamp of particle insertion or removal makes fixed-periodic tree rebuilding impractical. Parcels thus performs smart local-branch rebuilds at the time where particle indices in the ordering tree change. Specific challenges are the change of particle coordinates (and therefore indices) in a chain of per-particle kernels, as well as the proper construction of an unambiguous hitlist of closest particles. In short, the first challenge is addressed with the design decision to evaluate all advective kernels prior to any interaction, thus preventing index changes amid a single-pass kernel evaluation. The second challenge is addressed by the decision of symmetric ordering, i.e. *M*_0_(*q*_*i*_) = *q*_*j*_ ⟺ *M*_0_(*q*_*j*_) = *q*_*i*_ with *M*_*y*_(*q*_*z*_) being *y*th closest neighbour of particle *q*_*z*_. Those conditions are mandatory to resolve for particle-particle interaction to work in the oceanographic setting.

This specialised kD-Tree, which manages particle association in the background, facilitates relatively fast adjacency queries of particles without excessive memory overhead. This, in terms, enables us to model interactive behaviour between homogeneous- and heterogeneous types of particles, such as the FAD’s and tuna “super-individuals” in this study. Additionally, the computational method would in principle also facilitate direct predator-prey behaviour through using particles for both the tuna and their prey. However, the modelling and evaluation of prey as a field-quantity reduces the computational demand to a tenable level. As the presented study interest is the tuna, their prey is just consumed and thus requires no ABM or tracing in itself. Future studies can exploit the technical interaction possibilities even further by modelling dedicated swarm intelligence and group-aware behaviour, which is beyond the scope of this study.

In terms of constraints, the method as designed so far still requires tree rebuilding after each particle integration step with a *O*(3 × *n* log *n*) complexity, where *n* represents the number of particles, hence imposing significant computational costs for massive particle sets (*n* > 10^5^). In a parallel- and distributed computing setup, the kD tree construction and management needs to be globally consistent and thus needs to remain infull on one individual (computing) node. Implementation-wise, the method currently does not support multi-processing or distributed computing. Related method extensions need to address the significant communication overhead at each integration step, which is unavoidable due to the non-local nature of the algorithm. Lastly, the flexible particle definition and its various possible coordinate systems in oceanography makes a fast, *C*/*C*^++^-style implementation infeasible, hence all interaction-related kernels are evaluated in a computational setting based on Python and SciPy [55] exclusively.

### 2.5. Ocean flow configurations

Our model framework was run in several configurations across different, idealised flow fields (Table 1). Although these flow fields are idealised, they contain specific properties of realistic flows. First, we use a configuration where the flow is given by a Random Walk (RW): 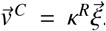. Here 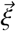 is a unit random vector, such that the value of *κ*^*R*^ (constant throughout the whole domain) determines the variance of the random walk. In this configuration, we use reflective boundary conditions for the particles. The initial prey field is uniformly distributed and with a value of 0.4. Depleted prey is redistributed at a uniformly random location. The RW flow represents an isotropically diffusive process.

Second, we use a Double Gyre flow [58], which we here refer to as the Double Eddy (DE) flow due to the scale at which it is applied (Fig. 1a). The Double Eddy flow has closed boundary conditions and we use reflective boundary conditions for the particles. Prey abundance is initialised as being large in the middle of one of the two eddies. The initialised prey field also provides the probability density function for the redistribution of depleted prey (depleted prey is more likely to be redistributed in the middle of this eddy). The DE flow has the property that passively advected particles accumulate in the middle of the two eddies over time, and hence represent meso-scale eddies that occur in the ocean.

**Figure 1:**
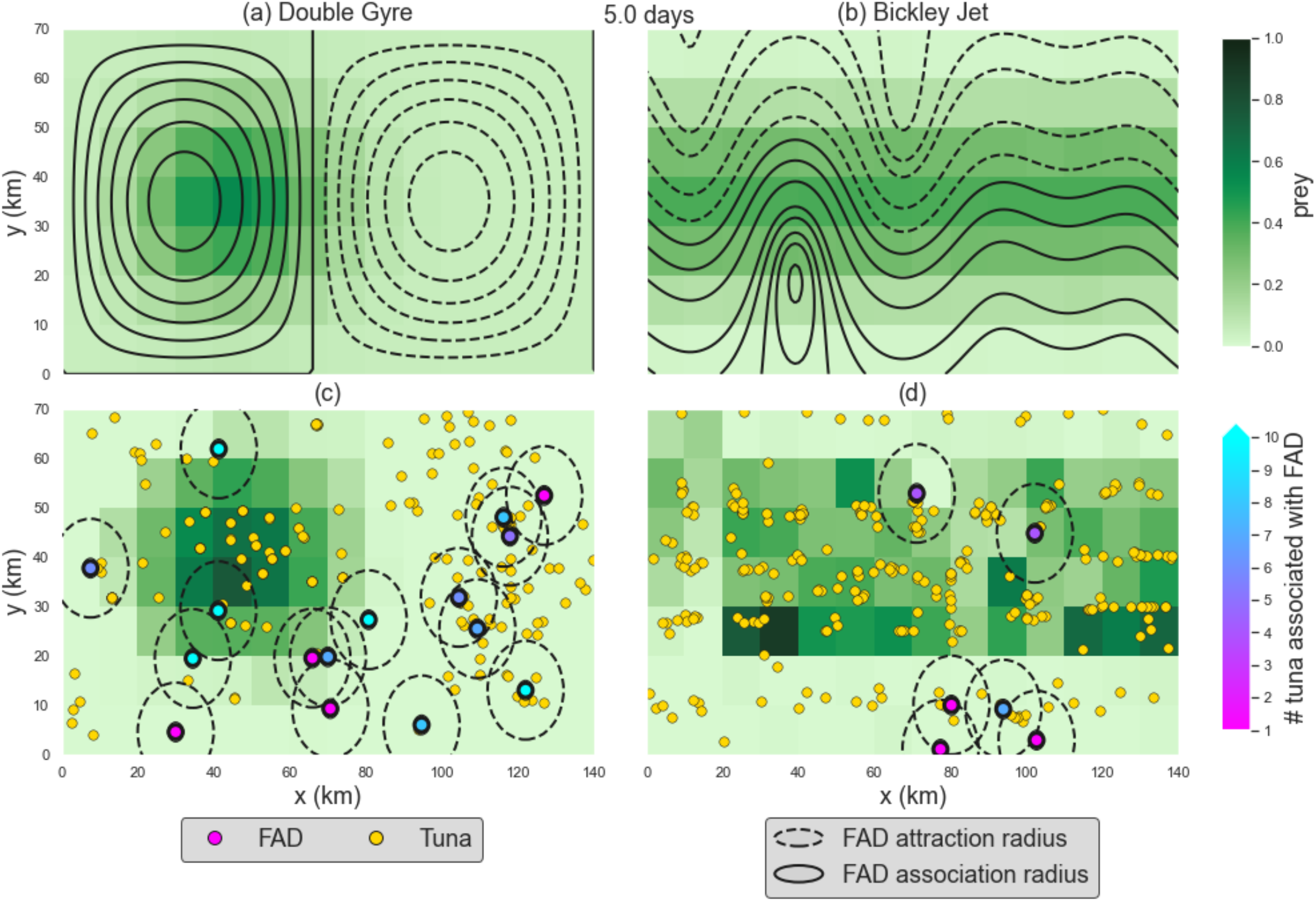
Streamlines of the (a) Double Eddy (DE) and (b) Bickley Jet (BJ) flow at *t*=5 days. Solid (dashed) lines represent positive (negative) values and (anti-)clockwise rotation. The background color represents the initialized prey field. Both flow fields are often used to study so-called Lagrangian coherent structures, that tuna tend to track in the real ocean [56, 57]. (c) and (d) snapshots at *t*=5 days of example simulations in the DE and BJ configurations, respectively. See the Supporting Information animations S1-S4 for the evolution of (a)-(d) in time.

Third, we use the Bickley Jet (BJ) flow [59] (Fig. 1b). At the spatial scale that is used here, the Bickley jet flow resembles an oceanic front. The Bickley jet has a zonal periodic boundary and a meridional closed boundary, which are also imposed as boundary conditions on the particles. The prey field is initialized to be large in the middle and low in the North and South of the domain.

The latter two flow configurations are often used when studying the dispersion of advected Lagrangian particles [60, 61]. The flow was scaled in these configurations, such that it fits on the fixed domain that we use in this paper, and the maximum flow velocity was set to the *κ*^*R*^ value of the random walk configuration. The code that was used to create these currents can be found on github (https://github.com/OceanParcels/InteractiveTuna).

In every configuration, the average prey index per grid box is set to the value *P*_*avg*_ = 0.1, the simulations are run for 100 days, *κ*^*I*^ = [0.01] and one tuna density (*N* = 500). Hence, we do not consider any tuna-FAD dynamics that may act on longer timescales than 100 days. See Supporting Information table S1 for other parameter values that are fixed in this paper. We will present the sensitivity of simulations on four different types of parameters. First, we will test the effect of tuna behaviour: *κ*^*T*^ = [0, 0.01], *κ*^*F*^ = [0, 0.5, 1] and *κ*^*P*^ = [0, 0.5, 1]. For clarity we summarise those parameter values under four behaviour hypotheses: (1) *κ*^*F*^ = *κ*^*P*^ and *κ*^*F*^; *κ*^*P*^ > 0 (i.e. FAD and prey attraction behaviour are equal), (2) *κ*^*F*^ > *κ*^*P*^ and *κ*^*F*^; *κ*^*P*^ > 0 (i.e. FAD-dominant attraction behaviour), (3) *κ*^*F*^ < *κ*^*P*^ and *κ*^*F*^; *κ*^*P*^ > 0 (i.e. prey dominant attraction behaviour), (4) *κ*^*F*^ = 0, or *κ*^*P*^ = 0 (i.e. one attraction behaviour switched off). Second, we test the four fishing strategies (i.e. FS0, FS1, FS2, FS3). Third, we test the outcomes for different FAD densities *F* =[0, 2, 5, 10, 15, 20, 30, 40]. Fourth, we compare differences between the RW, DE and BJ configurations. The total number of simulations presented in this paper is 1728.

### 2.6. Data

We present the results of our simulations across a suite of metrics, ranging from the internal state of individuals to the emergent dynamics of aggregated individuals within the domain. The patterns we compare our simulations with are: First, relative trends in numbers of tuna accumulated at FADs over time, prior to fishing events and thereafter, compared to observations of tuna biomass made by echo-sounder equipped FADs [5]. Second, the fullness of individuals’ stomachs, comparing to those from tuna caught and examined in the western central Pacific Ocean [62]. Third, the length of time that tuna spend associated with FADs, comparing to electronic tagging experiments of tuna around FADs [37, 21]. We have chosen these metrics so as to permit comparison to observed patterns and test the configurations of our model at multiple ecological scales, both spatiotemporal and hierarchical, simultaneously. Such an approach helps to constrain the high degrees of freedom inherent in individual-based models, for example by eliminating model configurations or structures that fail to produce the observed properties of a system at all levels [30]. Since we applied the ABM only in idealised and not in realistic configurations, we cannot set specific criteria on model performance. Therefore, we only compare whether these measures are of similar order of magnitude and represent similar patterns compared to observations.

## 3. Results

### 3.1. Colonisation of FADs by tuna

Unsurprisingly, more tuna become associated at FADs when tuna behaviour was FAD dominant compared to prey dominant (Fig. 2), since tuna are less likely to move away from FADs to forage. Tuna-tuna attraction had little influence on these dynamics (Supporting Information Fig. S1).

**Figure 2:**
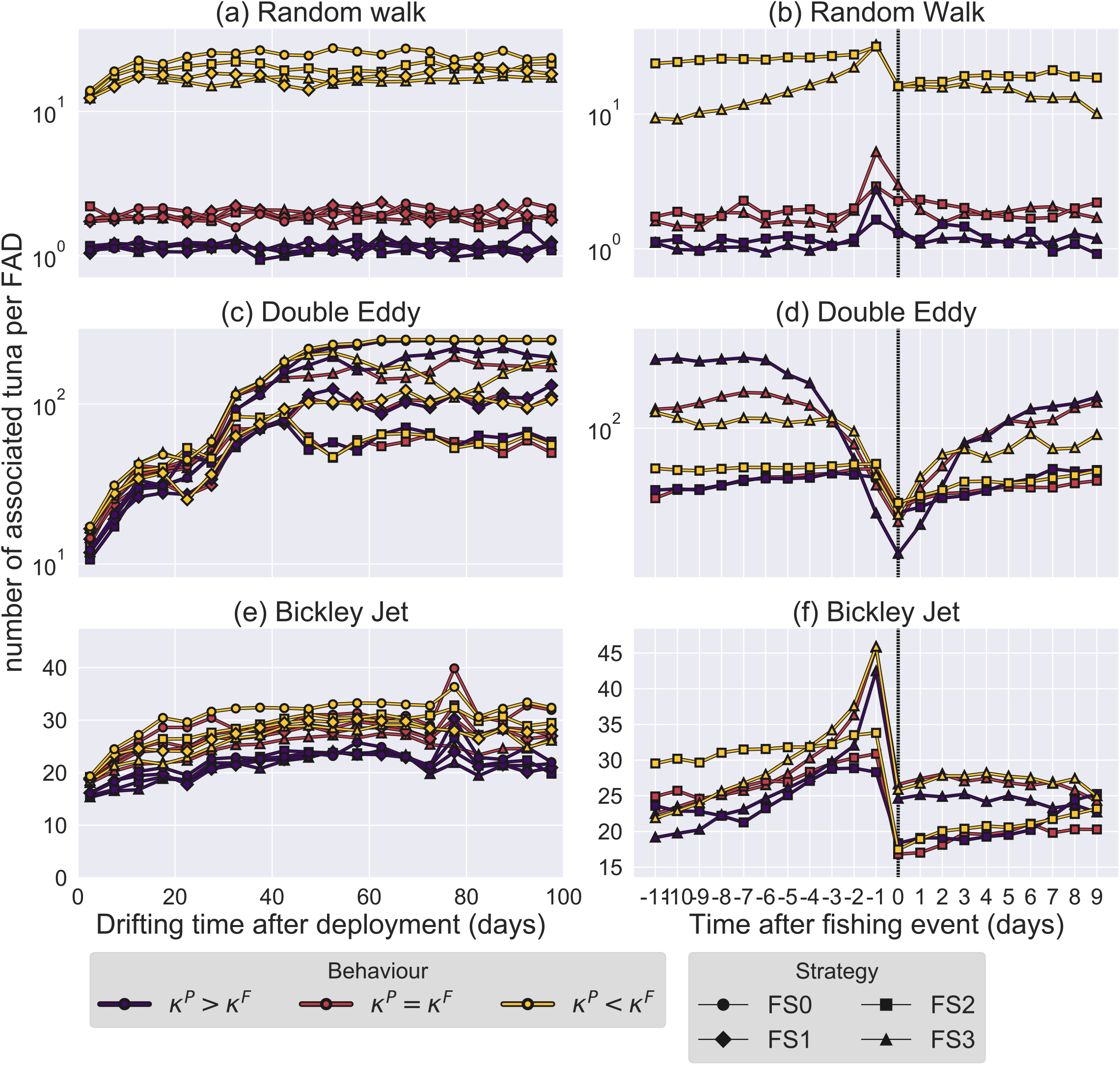
Colonisation of tuna at FADs after FAD deployment (left) and in the days before and after a fishing event (right; day 0 of fishing event is indicated with vertical dashed line). Median values of the number of associated tuna particles at FADs are shown for the (a), (b) Random Walk (c), (d) Double Eddy and (e), (f) Bickley Jet flow, averaged over parameter values with FAD (*κ*^*F*^ > *κ*^*P*^), prey (*κ*^*F*^ < *κ*^*P*^) and without dominant tuna behaviour and for different fishing strategies. Notice the logarithmic axis in (a)-(d). The FAD is density is *F* = 15 and tuna-tuna attraction is included (*κ*^*T*^ = 0.01; see Supporting Information Fig. S1 for the same figure with *κ*^*T*^ = 0).

Fishing strategies had a major influence on the distribution of tuna particles among FADs, as it governs how fish are depleted and redistributed throughout the domain. For a given behavioural configuration, fishing strategies that catch tuna more efficiently (e.g. FS3 compared to other fishing strategies), generally reduce the number of associated tuna at FADs during their initial colonisation (Fig. 2a, c, e). In the days leading up to fishing events, there was generally a larger number of associated tuna at the targeted FAD under fishing strategy FS3 compared to FS2 under the RW and BJ configurations (Fig. 2b, f), due to its more consistent targeting of large associations. This strategy prevents a buildup of large tuna numbers at several FADs at the same time. Before the number of associated tuna can grow substantially, a fishing event takes place at the FAD, resulting in a sudden halving of the number of associated tuna. In the case of the DE flow configuration, where both tuna and FAD particles tend to accumulate in the centre of the eddies, this can result in many tuna associated at FADs close to each other. Such associations can be repeatedly targeted for fishing under FS3, which results in an apparent decrease of biomass through time prior to fishing (see below), when averaged across the duration of a simulation. The distribution of tuna among FADs does not change much in the days after a fishing event, in part driven by the reduction in density-dependent, negative feedback from prey depletion (Fig. 2d,f), of which the strength is partly controlled by the relative *κ*^*P*^ and *κ*^*F*^ values.

Colonization dynamics only weakly depend on FAD density (Supporting Information Fig. S2). On average, it takes longer for tuna particles to get near a FAD if the FAD density is relatively low. As a result, it also takes longer for the colonization of tuna at FADs to stabilize after their deployment. Moreover, at relatively low FAD densities, less instances of large aggregations switch from one FAD to another prior to fishing, resulting in a smaller buildup of tuna particles at FADs before a fishing event.

FAD colonization and association dynamics differed strongly between different flow configurations (Fig. 2). In the RW configuration, tuna associate with nearby FADs in a few days after deployment (Fig. 2a). The distribution of the number of associated tuna at FADs does not change afterwards, even if large associations could be expected to build up under FS0 where no fishing events occur. This implies that tuna swim towards FADs if their stomach is full, where they subsequently deplete the local prey field. When the prey field gets depleted near the FAD due to the presence of many tuna particles and their stomach becomes empty, tuna start foraging and move away from the FAD. While this process repeats, the average number of tuna that associate with the FADs does not change, as the distribution of FADs is governed by a Brownian motion-like random walk. If tuna behaviour is FAD dominant, the build up of tuna at FADs takes somewhat longer (∼ 10 days) since tuna are less likely to swim away from a FAD. If tuna behaviour is prey dominant, few tuna associate with FADs, as a nonzero gradient of prey density can be easily found throughout the domain.

For the directed flow configurations DE and BJ, FADs are more likely to meet tuna particles compared to the RW flow. Under the DE flow, FADs are rapidly advected through either the westward or eastward eddy and tuna are attracted towards the prey rich area in the westward eddy where part of the FADs accumulate. Numbers of FAD associated tuna increase after deployment and stabilize after 50-80 days (Fig. 2c), when the FADs accumulate in the middle of the two eddies. On average, the build-up of tuna at FADs does not occur before a fishing event in the DE configuration (Fig. 2d), because the FADs accumulate in the middle of the two gyres and fishing events are very likely to occur at the same FADs every few days, which results in little associated tuna the day before these catch events.

The BJ domain-averaged flow has a clear eastward component. Similarly to the RW configuration, a rapid build up of associated tuna occurs in the first few days after deployment (Fig. 2e). However, the distribution is still changing until day 60, after which it stabilizes.

Observations from echo-sounder biomass estimates indicate that tuna colonisation at FADs after their deployment follows a log-normal distribution, where the amount of FAD associated tuna increases up to approximately 60-80 days after deployment (Fig. 12 in [5]), after which it stabilizes. This number reduces again on a time scale longer than the 100 day simulations presented in this paper. Both the BJ and DE configurations (Fig. 2c, e) compare better with these observations of tuna accumulation around FADs compared to the RW configuration, because the flow has a clear direction in BJ and DE, which is also often the case in reality. In the RW case on the other hand, the time-mean flow direction is 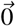 everywhere in the domain. Moreover, observations indicate a buildup of tuna at FADs before a fishing event and a stable number of associated tuna afterwards (Fig. 13 in [5]), similar to the RW and BJ configurations (Fig. 2b, f). Overall, we find that the ocean flow configuration has a major influence on the colonisation of tuna particles at FADs, chiefly due to FADs covering more distance over a given time within the domain, which aids in their ‘collection’ of tuna.

### 3.2. Stomach fullness

Tuna behaviour has implications for simulated stomach fullness. As more tuna particles become associated with a FAD due to FAD dominant behaviour, they deplete the prey locally (Fig. 3). As a consequence, the competition among tuna for food near them results in a lower stomach fullness of FAD associated tuna compared to unassociated tuna. When tuna particle swimming direction is only determined by the prey field (*κ*^*F*^ = 0), tuna either associate with FADs incidentally if the FAD is located in the prey rich area (e.g. western eddy in the DE flow), or if the flow field is responsible for the accumulation of both FAD and tuna particles in the same area (if no prey is available; e.g. in the eastern eddy in the DE flow). Unsurprisingly, stomach fullness is high if the tuna behaviour is not influenced by FADs (*κ*^*F*^ = 0), indicating that tuna particles are more likely to end up in prey rich areas when their swimming behaviour is only determined by prey. Although absolute stomach fullness does not change much when this is the case (*κ*^*F*^ = 0 compared to *κ*^*F*^ > 0), almost no difference exists between FAD associated and unassociated tuna. In contrast, when tuna swimming behaviour is only determined by FADs (*κ*^*P*^ = 0), those tuna that associate with a FAD, stay near the FAD until they are caught at a fishing event.

**Figure 3:**
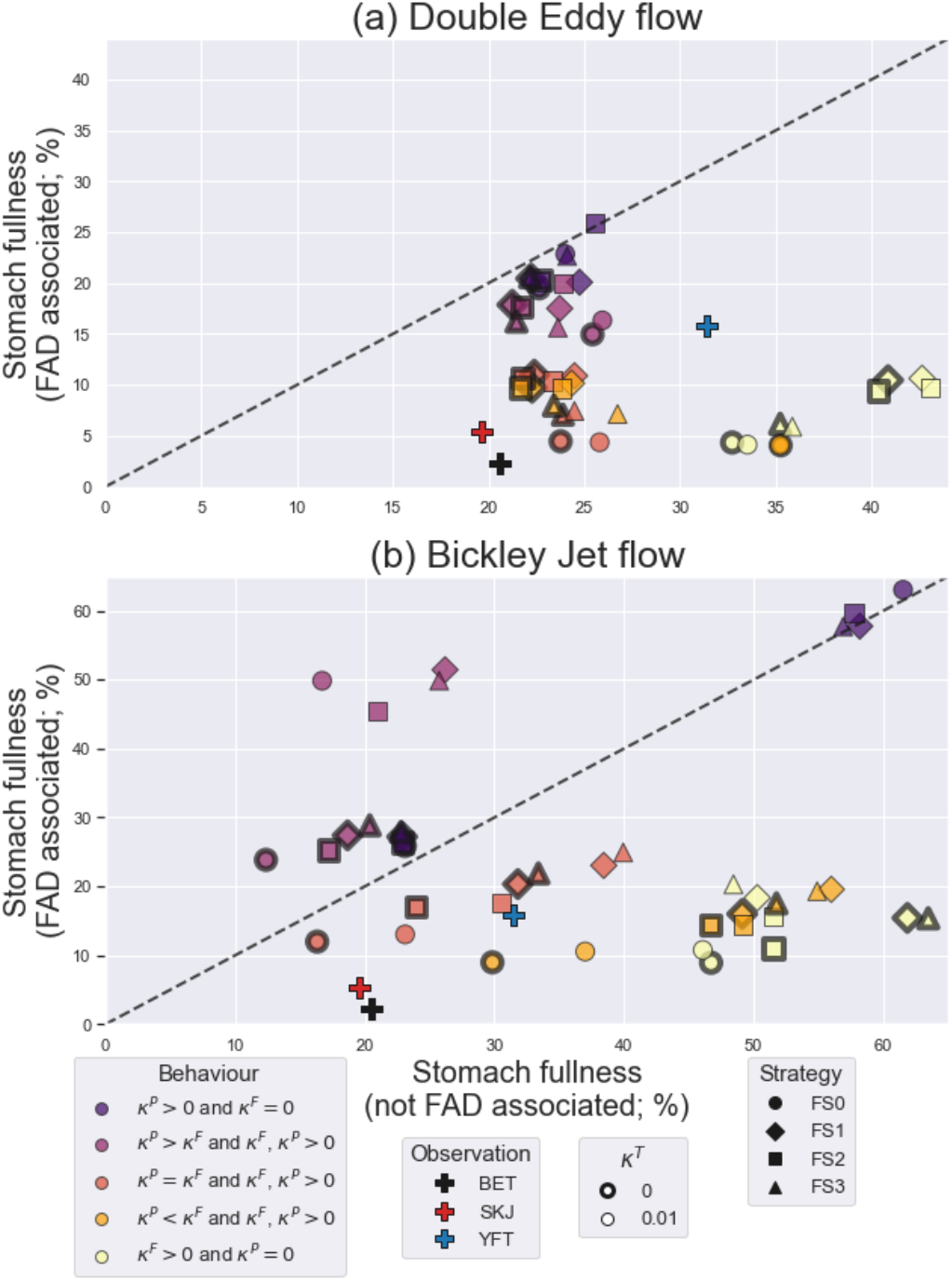
Tuna stomach fullness: associated versus not associated with FAD for the (a) Double Eddy (DE) and (b) Bickley Jet (BJ) flow configurations. Tuna stomach fullness is averaged over time, tuna particles and over parameter values with a specific tuna behaviour, for several fishing strategies and *κ*^*T*^ values. The FAD density is *F*=30. Observed stomach fullness [62] for bigeye (BET), skipjack (SKJ) and yellowfin (YFT) tuna are the same in (a) and (b). The black dashed line shows the one-to-one comparison where stomach fullness is the same for FAD associated and non-associated tuna. See Supporting Information Fig. S3 for the same figure with FAD density *F* = 5.

Tuna-tuna attraction increases tuna stomach fullness (*κ*^*T*^ = 0.01 versus *κ*^*T*^ = 0; on average a difference of ∼ 0.02 and ∼ 0.20 for DE and BJ configurations, respectively). Tuna-tuna attraction may lead to a ‘snowballing’ effect [6], where tuna may end up near a prey rich area by following other tuna. For instance, if tuna particles are located in the north or south of the domain in the BJ configuration, in the absence of any prey gradient or FADs, their swimming direction is arbitrary for *κ*^*T*^ = 0, and they are likely to remain here with an empty stomach. If *κ*^*T*^ > 0 on the other hand, it increases their ability to move to a prey rich area in the middle of the domain by following other tuna. As a result, tuna-tuna attraction results in less tuna particles near the northern or southern boundary of the domain, where prey abundance is low. Hence, tuna stomach fullness is generally higher if tuna-tuna attraction is included.

While FAD density has no clear effect on the stomach fullness (Supporting Information Fig. S3), the influence of fishing strategies on stomach fullness is only weak: on average tuna stomach fullness is ∼ 0.02 and ∼ 0.14 lower for fishing strategy FS0 compared to other fishing strategies for the DE and BJ configuration, respectively. Fishing strategies determine the amount of depletion and subsequent redistribution of tuna particles. More efficient fishing strategies imply that more tuna are caught and redistributed at a location without the presence of a FAD, which relieves the tuna density-dependent reduction of prey near the FAD. As a result, stomach fullness is generally higher for these efficient fishing strategies.

In the BJ flow configuration, stomach fullness is generally higher and its difference between FAD associated and non-associated tuna is more sensitive to tuna behaviour compared to the DE configuration (Fig. 3). The dominant flow direction in this configuration also helps to continuously transport FAD-associated tuna through the richest areas of the prey field, reducing local depletion and resulting in generally higher stomach fullness. We also tested the tuna stomach fullness in the RW flow configuration (not shown). In this configuration, stomach fullness is generally low (∼3%). Any tuna swimming behaviour results in a heterogeneous distribution of tuna particles, which focuses the depletion of the homogeneously distributed prey field. As a result, tuna particles are more often located in areas where the prey field is locally depleted. Since no dominant flow direction exists, this situation does not change. Hence, the tuna stomachs are generally low in this configuration.

Observations of tuna stomach contents indicate a lower stomach fullness for tuna that are caught near a FAD compared to tuna that are caught while not associated to a FAD (Fig. 3; ∼15%) [10, 63, 64, 62]. Note that it is mainly the comparison of the relative difference between FAD associated and non-associated tuna stomach fullness that is of importance here, since the observed stomach fullness itself is an indirect measure, varying with a number of covariates, and difficult to compare with the stomach fullness in our simulations. Both absolute and relative stomach fullness match well with observations in both the DE and BJ configuration (Fig. 3), especially for equal FAD- and prey-driven behaviour (*κ*^*P*^ = *κ*^*F*^).

### 3.3. Continuous Residence Times

Tuna leave a FAD either when they are caught, when another FAD pulls them away, or when they are foraging and move away towards an area with high prey abundance. Hence, we find that stronger FAD dominant behaviour increases Continuous Residence Times (CRT) and prey domainant behaviour decreases CRT, since the former increases the attraction strength of the FADs, and the latter causes abandonment of the FAD to hunt prey.

The impact of tuna-tuna attraction (*κ*^*T*^ > 0) on CRT is weak. Tuna-tuna attraction only has a relevant influence on CRT under fishing strategy FS0 (no fishing) in the BJ configuration (Fig. 4d). For FS0, where no tuna is caught and redistributed, the median CRT depends on how many tuna particles end up in the middle of the domain at the beginning of the simulation where prey abundance is high and cause the tuna to leave FADs while foraging. Tuna-tuna attraction may result in a higher tuna abundance in the middle of the domain through a ‘snowballing’ effect, and subsequent greater depletion of local prey [6].

**Figure 4:**
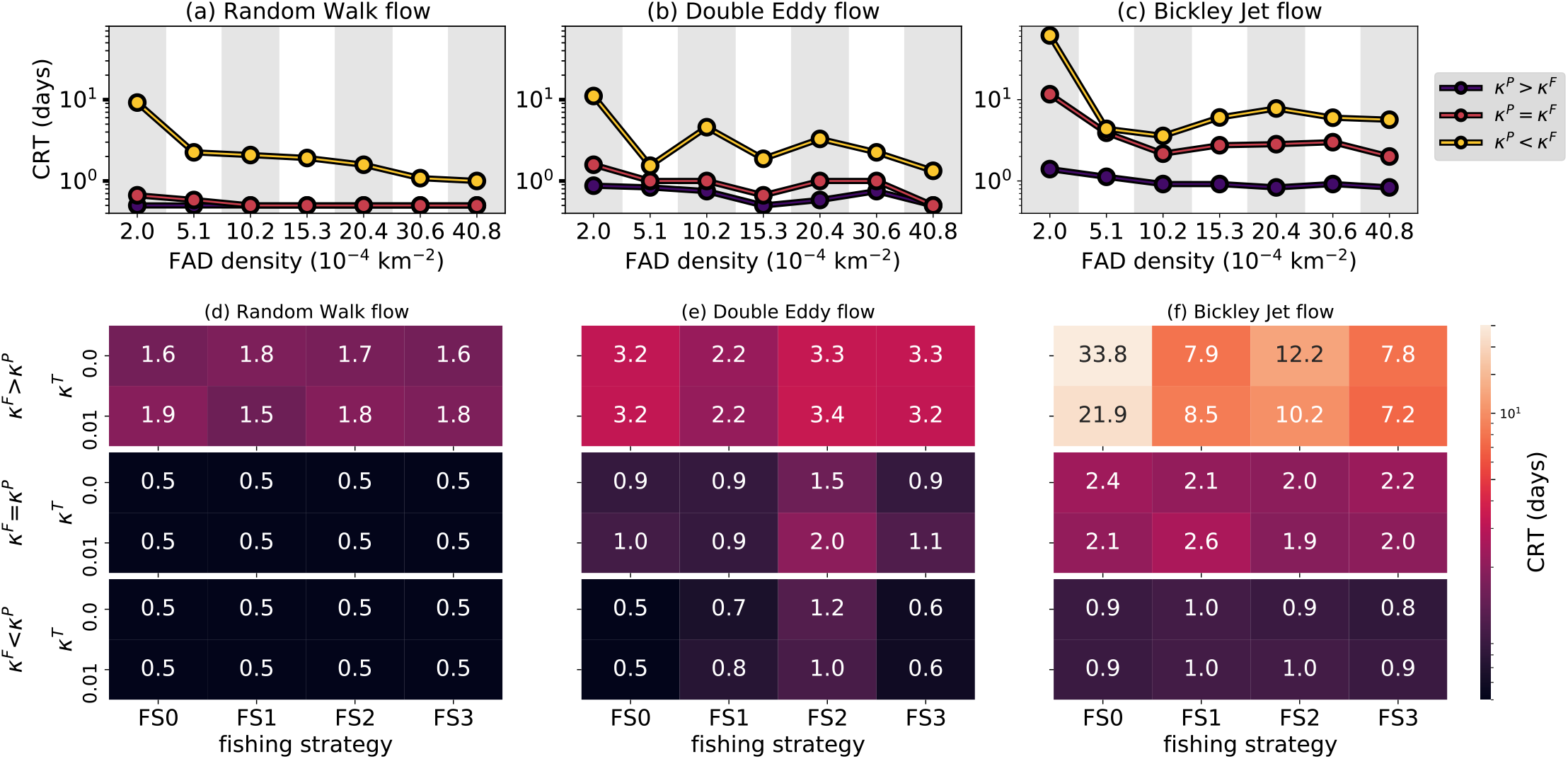
Median of the tuna Continuous Residence Times (CRT) at FADs in simulations with the Random Walk (left) Double Eddy (middle) and Bickley Jet (right) flow, averaged over parameter values with FAD (*κ*^*F*^ > *κ*^*P*^), prey (*κ*^*F*^ < *κ*^*P*^) and without (*κ*^*F*^ = *κ*^*P*^) dominant behaviour. (a)-(c) Median CRT at different FAD densities (fishing strategy FS2 and *κ*^*T*^ =0; notice the logarithmic vertical axis). (d)-(f) Median CRT for different fishing strategies and values of *κ*^*T*^ (FAD density *F*=20). Notice the logarithmic color scale.

Fishing strategies decrease CRT in our simulations, as removal of individuals from around FADs necessarily cuts short their residence at that FAD (Fig. 4d-f). However, this effect is distinctly different between ocean flow configurations. In the case of the BJ configuration, median CRT can be relatively long in the absence of fishing (FS0). CRTs are shorter in FS3 compared to FS2, because fishing is more efficient in FS3 and more fish are caught, which shortens the median CRT. Fishing strategies FS1 and FS2 do not deviate much, because clustering of tuna mostly occurs near FADs, and hence the fishing events also often occur near FADs under both strategies. There appeared to be little impact of fishing strategy on CRTs under the DE configuration, where resident times remained very short due to the accumulation of tuna and FADs in small areas, driving frequent switching between FADs by tuna (Supporting Information Fig. S4).

CRT decreases at higher FAD densities (Fig. 4a-c), in particular when tuna behaviour is either prey or FAD dominant (*κ*^*F*^ ≠ *κ*^*P*^). This pattern occurs because the increasing probability that groups of tuna particles move from one FAD to another. This switching of tuna from one FAD to another can occur either (a) because the other FAD, which has more associated tuna, is located nearby and is more attractive or (b) the prey field is locally depleted near the current FAD and tuna decide to look for prey elsewhere due to a low stomach fullness, and associated with a new FAD in a nearby and more prey rich area.

CRTs varied by flow field, being the longest in the BJ configuration compared to the RW and DE configurations, where the flow has a clear eastward direction, keeping tuna associated while continually moving into prey rich areas. As a consequence, tuna are unlikely to move away from a FAD to forage, as they move into prey rich areas while remaining associated with a FAD. RW is the configuration with shortest CRT, because tuna are most often forced to forage away from FADs in this configuration.

Median absolute CRT of tuna at FADs are mostly between 1-10 days in the simulations (Fig. 4), but can be several tens of days in some configurations (e.g. Fig. 4c, f), which conforms to observations [21, 37, 65] (Supporting Information Fig. S5). Observations in drifting (Supporting Information Fig. S5) and coastal, anchored FAD arrays [21] show an increasing CRT at higher FAD densities. In our simulations, increasing CRT for higher FAD densities only occurs for a few FAD densities in the DE and BJ configurations (Fig. 4b,c), although these do broadly match with FAD densities present in CRT observations. CRT can increase with FAD density, first because more FADs cover a larger part of the domain and will associate more tuna particles. Second, the same number of fishing events is distributed among more FADs. Hence, the probability reduces that a fishing event occurs at a specific FAD if the FAD density is higher, and the CRT of tuna associated with this FAD increases.

### 3.4. Tuna catch

We find that tuna catch is also sensitive to different behavioural parameters, flow configurations and fishing strategies in our simulations (Fig. 5). Catch is generally higher in those configurations that result in a more heterogeneous distribution of tuna particles, as our fishing strategies generally target aggregations. This is particularly the case for FAD dominant tuna behaviour (Fig. 5). Hence, FAD dominant behaviour results in a higher catch compared to prey dominant behaviour, if it adds to the heterogeneity of the tuna distribution (especially for the RW flow; Fig. 5d). However, tuna-tuna attraction influenced catch only weakly.

**Figure 5:**
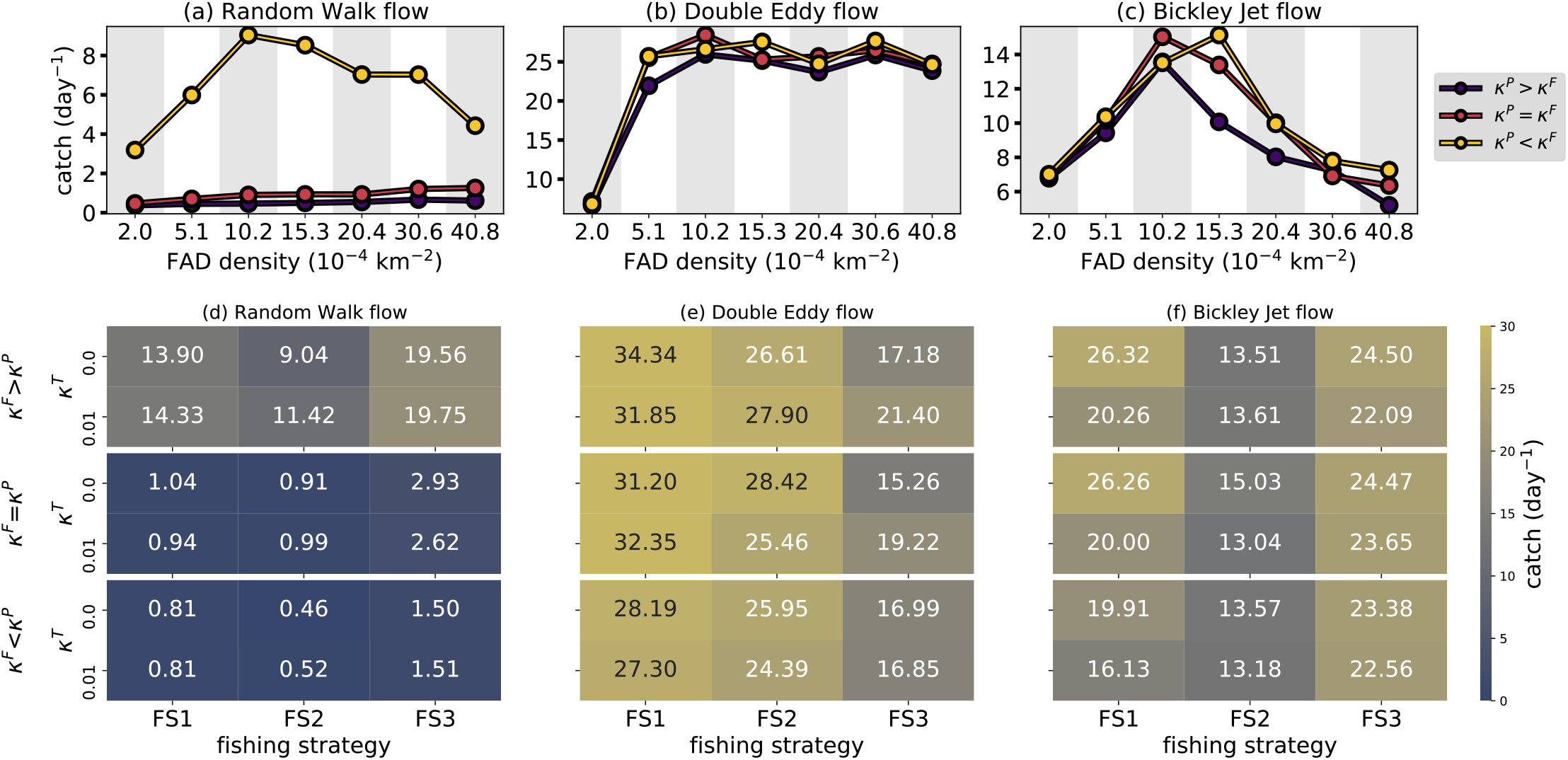
Average catch of tuna particles per day in simulations with the Random Walk (left), Double Eddy (middle) and Bickley Jet (right) flow, averaged over parameter values with FAD (*κ*^*F*^ > *κ*^*P*^), prey (*κ*^*F*^ < *κ*^*P*^) and without (*κ*^*F*^ = *κ*^*P*^) dominant behaviour. (a), (b), (c) Catch at different FAD densities and *κ*^*F*^, *κ*^*P*^ values (fishing strategy FS2 and *κ*^*T*^ =0). (d), (e), (f) Catch for different fishing strategies and values of *κ*^*T*^ (FAD density *F*=10).

The effect of fishing strategy varies markedly with ocean flow configuration. Within the BJ and RW flow, targeting of FADs with many associated tuna (FS3) always results in higher catch than targeting of random FADs (FS2). However, this relationship is reversed in the DE simulations (Fig. 5e), due to the concentration of particles in the centre of the eddies, and the density-dependant attraction towards FADs with many associated tuna becomes very strong. Since, the FS3 strategy reduces the number of associated tuna at densely populated FADs before assocations can grow to a substantial number, the attraction strength of FADs is reduced compared to the attraction strength of the prey field. As a result, mean catch actually decreases compared to evenly targeting FADs throughout the domain.

Interestingly, the targeting of random tuna particles in the domain (FS1) often results in a higher catch than random FAD fishing (FS2). Since FS1 picks a tuna particle at random to determine the fishing location, when tuna are heterogeneously distributed, it is likely that the fishing event occurs in an area with a lot of aggregated tuna, independently of whether those fish are associated with a FAD or not.

An optimal FAD density often exists where catch is maximized under the FS2 strategy (Fig. 5a, c). When FAD density is very low, only a fraction of the available tuna particles in the domain associate and are exposed to fishing, with the maximum possible aggregation limited by local depletion of prey. At very high FAD densities on the other hand, all available tuna particles are likely to associate with a FAD, but will be distributed over more FADs and the probability that a fishing event occurs on a large aggregation is reduced.

This optimum FAD density does not occur under the DE flow configuration. As this scenario results in very dense accumulation of FADs between which tuna switch frequently, targeting any FAD in the domain is likely to yield the same catch. As with our other results, the impact of tuna having generally emptier stomachs in the RW configuration reduces the time they associate and aggregate at FADs due to their increased need for foraging under greater competition, which subsequently reduces the total catch (Fig. 5a, d).

We do not compare the catch in our simulations with observations, because the number of tuna particles themselves cannot directly be compared to caught tuna biomass. Hence, if these simulations are compared to catch data, it can only be compared to the relative catch in different configurations. However, observed catch data represents many configurations of e.g. ocean flow, FAD density at the same time, and tuna catch is highly variable as a result. Moreover, general catch dynamics (e.g. the dependence of catch on FAD density as with Continuous Residence Times; CRT) are not known.

## 4. Discussion and outlook

Using the new particle-particle and particle-field interaction functionalities of the Parcels framework, we have developed an Agent-Based Model (ABM) of tuna behaviour. In this ABM, tuna is advected by ocean flow while interacting with drifting Fish Aggregating Devices (FADs), prey and other tuna.

The ABM presented here can be described by a small number of rules, but which allow for both direct and indirect interactions between components in the system to occur. The model was able to simultaneously produce many of the dynamics observed in the real ocean, such as the distribution of tuna among different FADs, their stomach fullness and the tuna continuous residence time near FADs. These emergent properties most consistently matched observations when simulated tuna behaviour was driven equally by both FADs and prey (*κ*^*F*^ = *κ*^*P*^).

However, testing the ABM in several idealised configurations of oceanic regime, we found that the flow and prey field configuration has a major impact on these emergent metrics. For instance, the dominant flow direction in the Bickley Jet scenario helps to reduce local prey depletion by large associated aggregations of tuna, allowing for a longer residence in the area around FADs without hunger causing dispersal due to foraging. Conversely, when there is an accumulation of particles due to flow as is the case in our Double Eddy scenario, concentration of both tuna and FADs occurs, with high levels of prey depletion and switching by aggregations in the centre of these eddies. Hence, the flow configuration determines the timescales at which tuna get close enough to a FAD to associate with it. In contrast, our simulations under a null, random-walk flow configuration, unrepresentative of the equatorial areas where tuna interact with FADs, resulted in far fewer of the real world patterns being replicated.

Great variability has been observed in the dynamics of tropical tunas around fish aggregating devices, causing suggestions of behavioural modes switching [24], high day-to-day variability in biomass detected by echosounder-equipped buoys [5], and mass abandonment of FADs by fish at the same time [11]. Here we have shown that such variation can occur at similar magnitudes purely as a function of changes to the flow field in which tuna and FADs find themselves, an oceanographic feature rarely incorporated into fisheries analyses [66].

The highly interactive nature of our agents mean that many of the emergent properties are highly sensitive to the fishing strategy. Beyond simply the depletion of tuna causing a reduction in the number of associated particles at a FAD, fishing events also relieve the density-dependent negative feedback of those associations, which deplete their surrounding prey field, whilst simultaneously reducing the positive feedback of FAD attraction caused by those large associations. This leads to counter-intuitive dynamics, such as targeting of tuna-associating with FADs for fishing actually increasing the mean residence time of fish, as it supports the remaining fish to stay associated for longer with lesser depletion of local prey around the FAD forcing a large association to fragment away from the FAD in search of food.

The ABM we have presented in this study remains idealised and with few parameters directly informed by data. However, such simulation models could be considered opaque thought-experiments [67], capable of exploring the potential mechanisms that lead to observed patterns and bracketing their uncertainty. Here we have shown that, for the set of biologically plausible assumptions we have structured our model on, realistic flow and prey fields are a requirement for observed dynamics to emerge. However, we also include a considerable flexibility in the behavioural parameters that can be used to examine more specific cases. Similarly, the real-world data that we have used for comparison of these dynamics have likely been observed across a range of ocean flow fields, tuna prey distributions, FAD densities and possibly even behavioural modes. Our ABM remains a hypothesis testing tool, where each of these components can be controlled. For example, tropical tuna behave differently among different species and size classes. Bigeye tuna (*Thunnus obesus*) associate with FADs during their small, juvenile stage, and slowly spend less time doing so as they grow and develop physiological adaptations to feed at depth. Although our ABM does not directly distinguish between different tuna types, several parameters (e.g. *κ*^*F*^, *κ*^*P*^) in the ABM determine the relative strength of different dynamics on the tuna behaviour. Hence, different of these parameter values may apply to these different classes or species, changing through time with the age of the fish.

We have chosen idealised flow and prey fields at the 1° scale to compare simulations across. This is typically the minimum scale at which tuna populations are modelled, and below which model processes are assumed to be homogeneous. Since these configurations are idealised, the distribution of tuna and their catch can be explained by known properties of the flow, prey, and interactions with FADs at this sub-grid scale. However, a question remains how these dynamics could be incorporated into ocean basin-scale tuna models [68]. For instance, these ocean basin-scale tuna models could use sub-grid scale parameterisations that are informed by similar ABM simulations of this paper. Identifying important sub-gride scale mechanisms, such as the impact of flow or FAD density on catch, then allows development of their incorporation into population-level models. This could be explored through alternate function responses, or as covariates in analyses that use catch per unit effort (CPUE) as an indication for population size. Furthermore, output from state-of-the-art hydrodynamic and biogeochemical models, or tuna species-specific habitat fields [68], could be incorporated to further test the ability of this or existing tuna ABMs [29] to replicate tuna-FAD dynamics in a more realistic scenario. In such an application, the prey field could be based on the underlying Eulerian model simulation and will be more compatible to the flow. Such a realistic setting may be applied to simulations in a future scenario, in order to test the influence of climate change on the distribution of FADs and its implication on their interaction with tuna. Applying the ABM in a realistic setting may require different fishing strategies. FS2 and FS3 are rather extreme cases where fishers either have no or almost complete knowledge on where the tuna is located. In reality however, their knowledge about the locations of tuna is likely somewhere in between fishing strategies FS2 and FS3.

The variability in simulated catch shown in this study highlights interesting questions regarding the relationship between catch, the density of FADs in a region, and whether certain model configurations will lead to a nonlinear response of catch to tuna abundance (so-called hyperstability or hyperdepletion). For example, the idealised scenarios we have explored in this study have shown that simulated catch from the same population size of tuna can change by a factor of two, purely as a result of different flow and prey field configurations, while all other parameters remain constant. Our tuna ABM could be used to investigate the dependence of catch levels on these parameters, across different tuna abundance under different model configurations. Examination of FAD-tuna observations, including catch, could where possible include in-situ oceanography to further test the hypotheses of our model. Such analyses would provide a better understanding of how to interpret catch data, potential levels at high stable populations may collapse, or parameterize tuna-FAD dynamics at a large-scale typically used to model their population dynamics and provide scientific advice on their management.

## Acknowledgments

The code used for this work is distributed under the MIT license and can be found at https://github.com/OceanParcels/InteractiveTuna. All authors thank the Information and Technology Service (ITS) of Utrecht University and specifically Roel Brouwer and Raoul Schram for their support with the particle-particle interaction development in Parcels. Funding was provided by the Western and Central Pacific Fisheries Commission (WCPFC Project 42) and the European Union “Pacific-European-Union-Marine-Partnership” Programme (agreement FED/2018/397-941). This publication was produced with the financial support of the European Union and Sweden. Its contents are the sole responsibility of the authors and do not necessarily reflect the views of the European Union and Sweden.

## Notes

https://github.com/OceanParcels/InteractiveTuna

## References

[1] R. Sharma, P. Levontin, T. Kitakado, L. Kell, I. Mosqueira, A. Kimoto, R. Scott, C. Minte-Vera, P. De Bruyn, Y. Ye, J. Kleineberg, J. L. Walton, S. Miller, A. Magnusson, Operating model design in tuna Regional Fishery Management Organizations: Current practice, issues and implications, Fish Fish. 21 (5) (2020) 940–961. doi:10.1111/faf.12480.

[2] A. Langley, A. Wright, G. Hurry, J. Hampton, T. Aqorua, L. Rodwell, Slow steps towards management of the world’s largest tuna fishery, Mar. Policy 33 (2) (2009) 271–279. doi:10.1016/j.marpol.2008.07.009.

[3] A. Maufroy, D. M. Kaplan, N. Bez, A. D. De Molina, H. Murua, L. Floch, E. Chassot, Massive increase in the use of drifting Fish Aggregating Devices (dFADs) by tropical tuna purse seine fisheries in the Atlantic and Indian oceans, ICES J. Mar. Sci. 74 (1) (2017) 215–225. doi:10.1093/icesjms/fsw175.

[4] A. Fonteneau, E. Chassot, N. Bodin, Global spatio-temporal patterns in tropical tuna purse seine fisheries on drifting fish aggregating devices (DFADs): Taking a historical perspective to inform current challenges, Aquat. Living Resour. 26 (1) (2013) 37–48. doi:10.1051/alr/2013046.

[5] L. Escalle, T. Vidal, B. Vanden Heuvel, R. Clarke, S. Hare, P. Hamer, G. Pillig, Project 88 final report: Acoustic FAD analyses, Tech. Rep. August, Western and Central Pacific Fisheries Commission (2021).

[6] J. J. Castro, J. A. Santiago, A. T. Santana-Ortega, A general theory on fish aggregation to floating objects: An alternative to the meeting point hypothesis, Rev. Fish Biol. Fish. 11 (3) (2002) 255–277. doi:10.1023/A:1020302414472.

[7] C. Girard, S. Benhamou, L. Dagorn, FAD: Fish Aggregating Device or Fish Attracting Device? A new analysis of yellowfin tuna movements around floating objects, Anim. Behav. 67 (2) (2004) 319–326. doi:10.1016/j.anbehav.2003.07.007.

[8] B. Leroy, J. Scutt Phillips, S. Nicol, G. M. Pilling, S. Harley, D. Bromhead, S. Hoyle, S. Caillot, V. Allain, J. Hampton, A critique of the ecosystem impacts of drifting and anchored FADs use by purse-seine tuna fisheries in the Western and Central Pacific Ocean, Aquat. Living Resour. 26 (1) (2013) 49–61. doi:10.1051/alr/2012033.

[9] L. Dagorn, E. Josse, P. Bach, A. Bertrand, Modeling tuna behaviour near floating objects: From individuals to aggregations, Aquat. Living Resour. 13 (4) (2000) 203–211. doi:10.1016/S0990-7440(00)01065-2.

[10] J. P. Hallier, D. Gaertner, Drifting fish aggregation devices could act as an ecological trap for tropical tuna species, Mar. Ecol. Prog. Ser. 353 (2008) 255–264. doi:10.3354/meps07180.

[11] G. Moreno, L. Dagorn, G. Sancho, D. Itano, Fish behaviour from fishers’ knowledge: The case study of tropical tuna around drifting fish aggregating devices (DFADs), Can. J. Fish. Aquat. Sci. 64 (11) (2007) 1517–1528. doi:10.1139/F07-113.

[12] P. Lehodey, I. Senina, R. Murtugudde, A spatial ecosystem and populations dynamics model (SEAPODYM) - Modeling of tuna and tuna-like populations, Prog. Oceanogr. 78 (4) (2008) 304–318. doi:10.1016/j.pocean.2008.06.004.

[13] J. Hampton, D. A. Fournier, A spatially disaggregated, length-based, age-structured population model of yellowfin tuna (Thunnus albacares) in the western and central pacific ocean, Mar. Freshw. Res. 52 (7) (2001) 937–963. doi:10.1071/MF01049.

[14] D. A. Fournier, J. Hampton, J. R. Sibert, MULTIFAN-CL: A length-based, age-structured model for fisheries stock assessment, with application to South Pacific albacore, Thunnus alalunga, Can. J. Fish. Aquat. Sci. 55 (9) (1998) 2105–2116. doi:10.1139/f98-100.

[15] C. M. Petrik, C. A. Stock, K. H. Andersen, P. D. van Denderen, J. R. Watson, Bottom-up drivers of global patterns of demersal, forage, and pelagic fishes, Prog. Oceanogr. 176 (May) (2019) 102124. doi:10.1016/j.pocean.2019.102124.

[16] K. A. Kearney, S. J. Bograd, E. Drenkard, F. A. Gomez, M. Haltuch, A. J. Hermann, M. G. Jacox, I. C. Kaplan, S. Koenigstein, J. Y. Luo, M. Masi, B. Muhling, M. Pozo Buil, P. A. Woodworth-Jefcoats, Using Global-Scale Earth System Models for Regional Fisheries Applications, Front. Mar. Sci. 8 (August) (2021) 1–27. doi:10.3389/fmars.2021.622206.

[17] A. J. Lotka, Elements of Physical Biology, Nature 116 (2917) (1925) 461.

[18] V. Volterra, Fluctuations in the Abundance of a Species considered Mathematically, Nature 118 (1926) 558–560.

[19] C. S. Holling, Some Characteristics of Simple Types of Predation and Parasitism, Can. Entomol. 91 (7) (1959) 385–398.

[20] F. Arregúin-Sánchez, Catchability: A key parameter for fish stock assessment, Rev. Fish Biol. Fish. 6 (2) (1996) 221–242. doi:10.1007/bf00182344.

[21] G. Pérez, L. Dagorn, J. L. Deneubourg, F. Forget, J. D. Filmalter, K. Holland, D. Itano, S. Adam, R. Jauharee, S. P. Beeharry, M. Capello, Effects of habitat modifications on the movement behavior of animals: the case study of Fish Aggregating Devices (FADs) and tropical tunas, Mov. Ecol. 8 (1) (2020) 1–10. doi:10.1186/s40462-020-00230-w.

[22] K. M. Schaefer, D. W. Fuller, Vertical movements, behavior, and habitat of bigeye tuna (thunnus obesus) in the equatorial eastern pacific ocean, ascertained from archival tag data, Marine Biology 157 (12) (2010) 2625–2642.

[23] J. Scutt Phillips, G. M. Pilling, B. Leroy, K. Evans, T. Usu, C. H. Lam, K. M. Schaefer, S. Nicol, Revisiting the vulnerability of juvenile bigeye (Thunnus obesus) and yellowfin (T. albacares) tuna caught by purse-seine fisheries while associating with surface waters and floating objects, PLoS One 12 (6) (2017) 1–18. doi:10.1371/journal.pone.0179045.

[24] M. Robert, L. Dagorn, J. D. Filmalter, J. L. Deneubourg, D. Itano, K. Holland, Intra-individual behavioral variability displayed by tuna at fish aggregating devices (FADs), Mar. Ecol. Prog. Ser. 484 (2013) 239–247. doi:10.3354/meps10303.

[25] Y. Baidai, M. Amande, D. Gaertner, L. Dagorn, M. Capello, Recent advances on the use of supervised learning algorithms for detecting tuna aggregations under fads from echosounder buoys data, Tech. rep., IOTC-2018-WPTT20-25 Rev1. Mahé (2018).

[26] D. Precioso, M. Navarro-García, K. Gavira-O’Neill, A. Torres-Barrán, D. Gordo, V. Gallego-Alcalá, D. Gómez-Ullate, Tuna-ai: tuna biomass estimation with machine learning models trained on oceanography and echosounder fad data, arXiv preprint 2109.06732 (2021).

[27] J. Lopez, G. Moreno, C. Lennert-Cody, M. Maunder, I. Sancristobal, A. Caballero, L. Dagorn, Environmental preferences of tuna and non-tuna species associated with drifting fish aggregating devices (dfads) in the atlantic ocean, ascertained through fishers’ echo-sounder buoys, Deep Sea Research Part II: Topical Studies in Oceanography 140 (2017) 127–138.

[28] L. Dagorn, P. Fréon, Tropical tuna associated with floating objects: A simulation study of the meeting point hypothesis, Can. J. Fish. Aquat. Sci. 56 (6) (1999) 984–993. doi:10.1139/cjfas-56-6-984.

[29] J. Scutt Phillips, A. Sen Gupta, I. Senina, E. van Sebille, M. Lange, P. Lehodey, J. Hampton, S. Nicol, An individual-based model of skipjack tuna (Katsuwonus pelamis) movement in the tropical Pacific ocean, Prog. Oceanogr. 164 (February) (2018) 63–74. doi:10.1016/j.pocean.2018.04.007.

[30] V. Grimm, E. Revilla, U. Berger, F. Jeltsch, W. M. Mooij, S. F. Railsback, H. H. Thulke, J. Weiner, T. Wiegand, D. L. DeAngelis, Pattern-oriented modeling of agent-based complex systems: Lessons from ecology, Science 310 (5750) (2005) 987–991. doi:10.1126/science.1116681.

[31] V. Grimm, S. F. Railsback, Pattern-oriented modelling: A ‘multi-scope’ for predictive systems ecology, Philos. Trans. R. Soc. B Biol. Sci. 367 (1586) (2012) 298–310. doi:10.1098/rstb.2011.0180.

[32] Y. Tyutyunov, L. Titova, R. Arditi, Predator interference emerging from trophotaxis in predator-prey systems: An individual-based approach, Ecol. Complex. 5 (1) (2008) 48–58. doi:10.1016/j.ecocom.2007.09.001.

[33] R. Arditi, Y. Tyutyunov, A. Morgulis, V. Govorukhin, I. Senina, Directed movement of predators and the emergence of density-dependence in predator-prey models, Theor. Popul. Biol. 59 (3) (2001) 207–221. doi:10.1006/tpbi.2001.1513.

[34] V. Castellanos, R. E. Chan-López, Existence of limit cycles in a three level trophic chain with Lotka–Volterra and Holling type II functional responses, Chaos, Solitons and Fractals 95 (2017) 157–167. doi:10.1016/j.chaos.2016.12.011.

[35] C. Kehl, D. Reijnders, R. Fisher, R. Brouwer, R. Schram, E. van Sebille, Parcels 2.2 - an increasingly versatile, open-source lagrangian ocean simulation tool, EGU General Assembly 2021, online, 19–30 Apr 2021 1033 (2021).

[36] D. S. Kirby, G. Allain, P. Lehodey, A. Langley, Individual/agent-based modelling of fishes, fishers, and turtles, in: 17 th Meeting of the Standing Committee on Tuna and Billfish, Majuro, Republic of Marshall Islands, 2004, pp. 9–18.

[37] J. Scutt Phillips, B. Leroy, T. Peatman, L. Escalle, N. Smith, Electronic tagging for the mitigation of bigeye and yellowfin tuna juveniles by purse seine fisheries, Tech. Rep. August, Western and Central Pacific Fisheries Commission (2019).

[38] M. Becher, V. Grimm, P. Thorbek, J. Horn, P. kennedy, J. Osborne, Beehave: a systems model of honeybee colony dynamics and foraging to explore multifactorial causes of colony failure, Journal of applied ecology 51 (2014) 470–482.

[39] B. Meyer, U. Freier, V. Grimm, J. Groeneveld, B. Hunt, S. Kerwath, R. King, C. Klaas, E. Pakhomov, K. Meiners, J. Melbourne-Thomas, E. Murphy, S. Thorpe, S. Stammerjohn, D. Wolf-Gladrow, L. Auerswald, A. Gotz, L. Halbach, S. Jarman, S. Kawaguchi, T. Krumpen, G. Nehrke, R. Ricker, M. Sumner, M. Teschke, R. Trebilco, N. Yilmaz, The winter pack-ice zone provides a sheltered but food-poor habitat for larval antarctic krill, Nature ecology and evolution 1 (2017) 1853–1861.

[40] R. B. Cabral, P. M. Alino, M. T. Lim, Modelling the impacts of fish aggregating devices (FADs) and fish enhancing devices (FEDs) and their implications for managing small-scale fishery, ICES J. Mar. Sci. 71 (7) (2014) 1750–1759. doi:10.1038/278097a0.

[41] M. Capello, M. Soria, P. Cotel, G. Potin, L. Dagorn, P. Fréon, The het-erogeneous spatial and temporal patterns of behavior of small pelagic fish in an array of Fish Aggregating Devices (FADs), J. Exp. Mar. Bio. Ecol. 430-431 (2012) 56–62. doi:10.1016/j.jembe.2012.06.022.

[42] D. Stephens, J. Krebs, Foraging theory, Princeton University Press, 1st edition, 1978.

[43] T. J. Pitcher, The behaviour of teleost fishes, Springer Science & Business Media, 2012.

[44] G. M. Viswanathan, M. G. Da Luz, E. P. Raposo, H. E. Stanley, The physics of foraging: an introduction to random searches and biological encounters, Cambridge University Press, 2011.

[45] A. Okubo, S. A. Levin, Diffusion and ecological problems: modern perspectives, Vol. 14, Springer, 2001.

[46] J. J. Magnuson, J. J. Magnuson, Digestion and Food Consumption by Skipjack Tuna (Katsuwonus pelamis) Digestion and Food Consumption by Skipjack Tuna (Katsuwonus pelamis), Trans. Am. Fish. Soc. 98 (3) (1969) 37–41.

[47] R. W. Brill, Selective advantages conferred by the high performance physiology of tunas, billfishes, and dolphin fish, Comp. Biochem. Physiol. -A Physiol. 113 (1) (1996) 3–15. doi:10.1016/0300-9629(95)02064-0.

[48] B. Faugeras, O. Maury, Modeling fish population movements: From an individual-based representation to an advection-diffusion equation, J. Theor. Biol. 247 (4) (2007) 837–848. doi:10.1016/j.jtbi.2007.04.012.

[49] M. Robert, L. Dagorn, J. L. Deneubourg, The aggregation of tuna around floating objects: What could be the underlying social mechanisms?, J. Theor. Biol. 359 (2014) 161–170. doi:10.1016/j.jtbi.2014.06.010.

[50] F. Tolentino-Zonderva, P. Berentsen, S. Bush, A. Oude Lansink, Fad vs. free school: Effort allocation by marine stewardship council compliant filipino tuna purse seiners in the pna, Marine Policy 90 (2018) 137–145.

[51] P. Delandmeter, E. van Sebille, The Parcels v2.0 Lagrangian framework: new field interpolation schemes, Geosci. Model Dev. 12 (2019) 3571–3584.

[52] M. Müller, B. Solenthaler, R. Keiser, M. Gross, Particle-based fluidfluid interaction, SCA ’05: Proceedings of the 2005 ACM SIGGRAPH/Eurographics symposium on Computer animation (2005) 237–244.

[53] M. Liu, G. Liu, Smoothed particle hydrodynamics (sph): an overview and recent developments, Archives of Computational Methods in Engineering volume 17 (2010) 25–76.

[54] L. Linsen, T. Van Long, P. Rosenthal, S. Rosswog, Surface extraction from multi-field particle volume data using multi-dimensional cluster visualization., IEEE transactions on visualization and computer graphics 14 (6) (2008) 1483–1490.

[55] P. Virtanen, R. Gommers, T. E. Oliphant, M. Haberland, T. Reddy, D. Cournapeau, E. Burovski, P. Peterson, W. Weckesser, J. Bright, S. J. van der Walt, M. Brett, J. Wilson, K. J. Millman, N. Mayorov, A. R. J. Nelson, E. Jones, R. Kern, E. Larson, C. J. Carey, İ. Polat, Y. Feng, E. W. Moore, J. VanderPlas, D. Laxalde, J. Perktold, R. Cimrman, I. Henriksen, E. A. Quintero, C. R. Harris, A. M. Archibald, A. H. Ribeiro, F. Pedregosa, P. van Mulbregt, SciPy 1.0 Contributors, SciPy 1.0: Fundamental Algorithms for Scientific Computing in Python, Nature Methods 17 (2020) 261–272. doi:10.1038/s41592-019-0686-2.

[56] S. V. Prants, M. V. Budyansky, M. Y. Uleysky, V. V. Kulik, Lagrangian fronts and saury catch locations in the Northwestern Pacific in 2004–2019, J. Mar. Syst. 222 (2021). doi:10.1016/j.jmarsys.2021.103605.

[57] E. T. Kai, V. Rossi, J. Sudre, H. Weimerskirch, C. Lopez, E. Hernandez-Garcia, F. Marsac, V. Garcon, Top marine predators track Lagrangian coherent structures, Proc. Natl. Acad. Sci. 106 (20) (2009) 8245–8250. doi:10.1073/pnas.0811034106.

[58] S. C. Shadden, F. Lekien, J. E. Marsden, Definition and properties of Lagrangian coherent structures from finite-time Lyapunov exponents in two-dimensional aperiodic flows, Phys. D Nonlinear Phenom. 212 (3-4) (2005) 271–304. doi:10.1016/j.physd.2005.10.007.

[59] W. Bickley, LXXIII. The plane jet, London, Edinburgh, Dublin Philos. Mag. J. Sci. 23 (1937) 727–731. doi:10.1080/14786443708561847.

[60] G. Conti, G. Badin, Hyperbolic covariant coherent structures in two dimensional flows, Fluids 2 (4) (2017). doi:10.3390/fluids2040050.

[61] D. Del-Castillo-Negrete, P. J. Morrison, Chaotic transport by Rossby waves in shear flow, Phys. Fluids A 5 (4) (1992) 948–965. doi:10.1063/1.858639.

[62] P. Machful, A. Portal, J. Macdonald, V. Allain, J. Scutt Phillips, S. Nicol, Tuna stomachs: Is the glass half full, or half empty?, SPC Fisheries Newsletter 166 (2021) 38–44.

[63] F. Ménard, B. Stéquert, A. Rubin, M. Herrera, É. Marchal, Food consumption of tuna in the equatorial atlantic ocean: Fad-associated versus unassociated schools, Aquatic living resources 13 (4) (2000) 233–240.

[64] V. Allain, B. Leroy, Ecosystem monitoring and analysis: stomach sampling overview of the gef-sap project 2000-2005 and stomach sampling strategy of the gef-ofm project, Tech. rep., Western and Central Pacific Fisheries Commission (2006).

[65] J. Scutt Phillips, G. M. Pilling, B. Leroy, K. Evans, T. Usu, C. H. Lam, K. M. Schaefer, S. Nicol, Revisiting the vulnerability of juvenile bigeye (thunnus obesus) and yellowfin (t. albacares) tuna caught by purse-seine fisheries while associating with surface waters and floating objects, PloS one 12 (6) (2017) e0179045.

[66] T. Vidal, P. Hamer, L. Escalle, G. Pilling, Assessing trends in skipjack tuna abundance from purse seine catch and effort data in the wcpo, Tech. rep., Technical Report WCPFC-SC16–2020/SA-IP-09 (2020).

[67] E. A. Di Paolo, J. Noble, S. Bullock, Simulation models as opaque thought experiments, in: M. A. Bedau, J. S. McCaskill, N. Packard, S. Rasmussen (Eds.), Artificial Life VII: Proceedings of the Seventh International Conference on Artificial Life, MIT Press, Cambridge, MA, 2000, pp. 497–506.

[68] I. Senina, P. Lehodey, J. Sibert, J. Hampton, Integrating tagging and fish1130 eries data into a spatial population dynamics model to improve its predictive skills, Canadian Journal of Fisheries and Aquatic Sciences 77 (3) (2020) 576–593.

