## Supplementary material for "Modelling of tuna around fish aggregating devices: the importance of ocean flow and prey": SupportingInformation.pdf

---

**Contents of this file**

1. Figures S1 to S5
2. Table S1

**Additional Supporting Information (Files uploaded separately)**

1. Captions for Movies S1 to S4

**Movie S1.** Animation of tuna and FAD particles in the Double Eddy flow (from Fig. 1c), $\kappa^F = 1$, $\kappa^P = 1$, $\kappa^T = 0$, fishing strategy FS2.

**Movie S2.** Animation of tuna and FAD particles in the Bickley Jet flow (from Fig. 1d), $\kappa^F = 1$, $\kappa^P = 1$, $\kappa^T = 0$, fishing strategy FS2.

**Movie S3.** Animation of tuna and FAD particles in the Random Walk flow, $\kappa^F = 1$, $\kappa^P = 1$, $\kappa^T = 0$, fishing strategy FS2.

**Movie S4.** Animation of the streamlines of the Double Eddy and the Bickley Jet flow (from Fig. 1a,b). Solid lines denote positive values and clockwise rotation. Dashed lines denote negative values and anti-clockwise rotation.

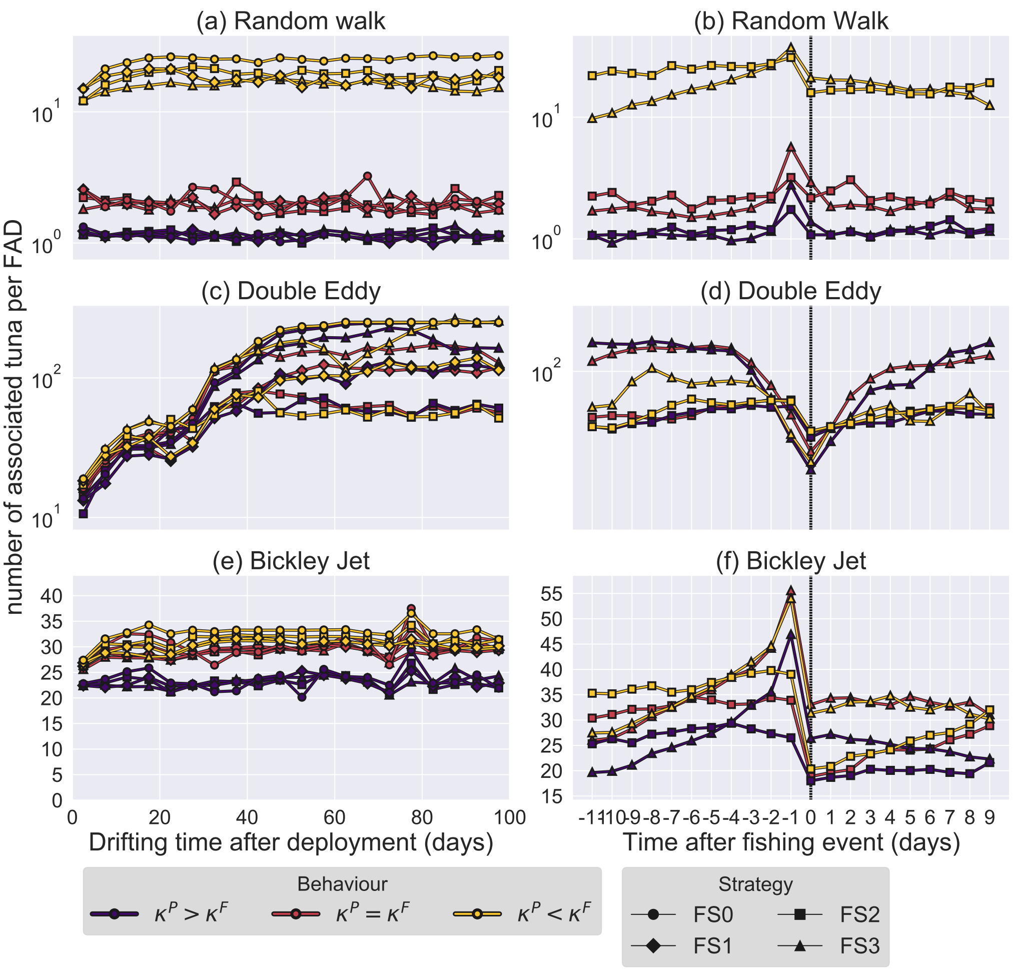


Figure S1: Same as Fig 2, but excluding tuna-tuna attraction ($\kappa^T = 0$).

Table S1: Overview of model parameters. Parameter values are only given if they are constant throughout the paper.

| parameter | value | description |
|---|---|---|
| $L_x$ | 140km | Zonal domain size |
| $L_y$ | 70km | Meridional domain size |
| $\Delta t$ | 20 minutes | Time step |
| $L^F, L^P$ | 1,2 | Determines the maximum of logistic function $l(n)$ |
| $k^F, k^P$ | 0.35, 10 | Steepness of logistic function $l(n)$ [1] |
| $n_0^F, n_0^P$ | 12, 0.5 | Midpoint of logistic function $l(n)$ [1] |
| $C^F, C^P$ | 1, 0 | Minimum of logistic function $l(n)$ [1] |
| $\epsilon^P \in [0, 1]$ | $1.4 \cdot 10^{-4}\text{s}^{-1}$ | The amount by which the prey field is depleted by tuna [1] |
| $\epsilon^E \in [0, 1]$ | $2.3 \cdot 10^{-5}\text{s}^{-1}$ | Tuna stomach Evacuation [2, 3] |
| $\epsilon^T$ | 0.5 | Catch probability at a fishing event |
| $\beta$ | 300 | Scale between tuna ingestion rate and field grid cell depletion |
| $\alpha$ | 3 | von Mises concentration parameter of tuna-prey attraction |
| $\gamma$ | 2 | von Mises concentration parameter of tuna-tuna attraction |
| $v_{max}$ | 0.4m $\text{s}^{-1}$ | Maximum swimming velocity of tuna [4, 5] |
| $R_a$ | 2km | FAD association radius |
| $R_b$ | 10km | FAD-tuna interaction distance [6, 7] |
| $R_c$ | 3km | Interaction distance between tuna particles |
| $\Delta x$ | 10km | Horizontal resolution of the prey field |
| $\kappa^R$ | 0.05m $\text{s}^{-1}$ | Determines ocean flow strength |
| $N$ | 500 | Total number of tuna particles |
| $P_{avg}$ | 0.1 | Mean prey index per grid cell |
| $\kappa^I$ | 0.01 | Relative contribution of inertia to tuna swimming direction [8] |
| $\kappa^F$ | | Relative contribution of FADs to tuna swimming direction |
| $\kappa^T$ | | Relative contribution of other tuna to tuna swimming direction |
| $\kappa^P$ | | Relative contribution of prey field to tuna swimming direction |
| $p$ | | Determines the geometric distribution shape for fishing strategies FS2 and FS3 |
| $F$ | | Number of FAD particles in the domain |

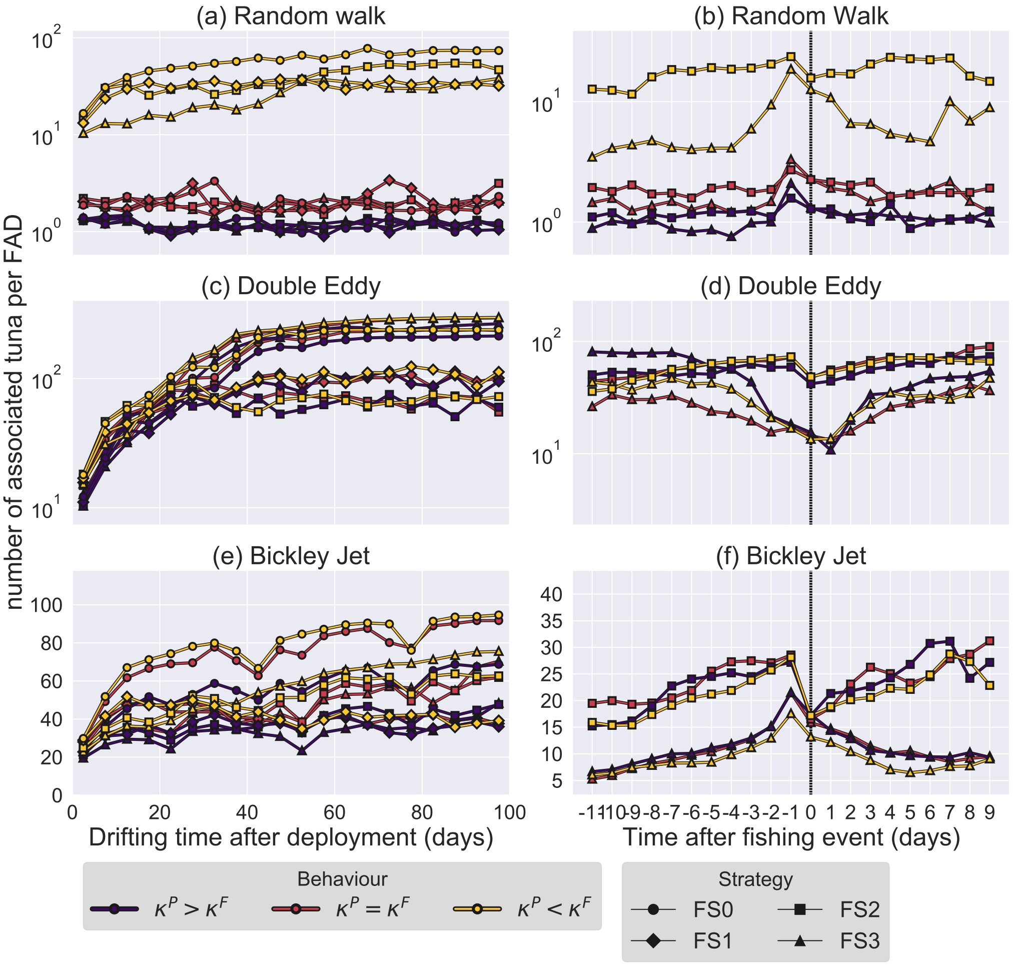


Figure S2: Same as Fig 2, but excluding tuna-tuna attraction ($\kappa^T = 0$) and FAD density $F = 5$.

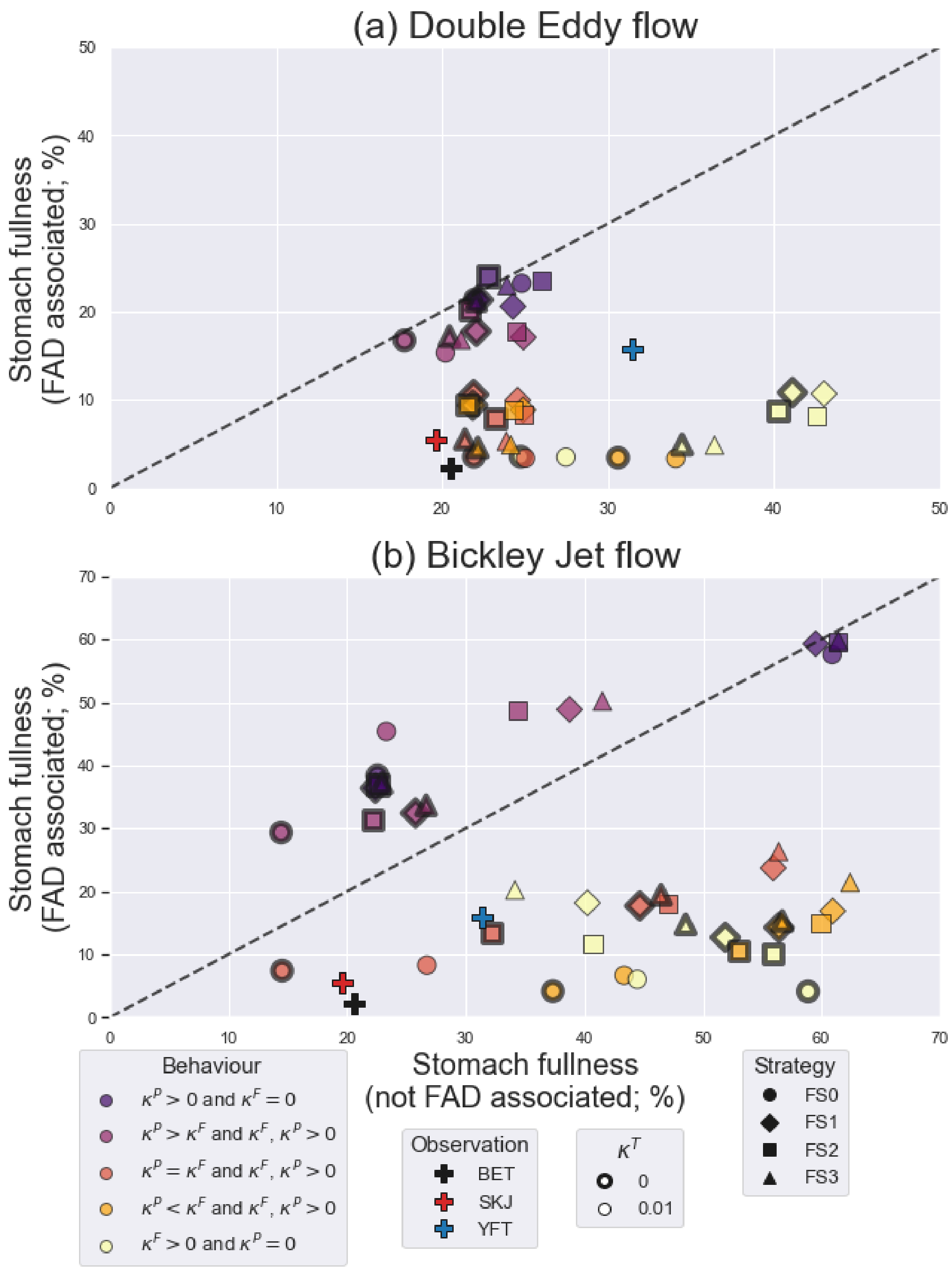


Figure S3: Same as Fig. 3, but with FAD density $F = 15$.

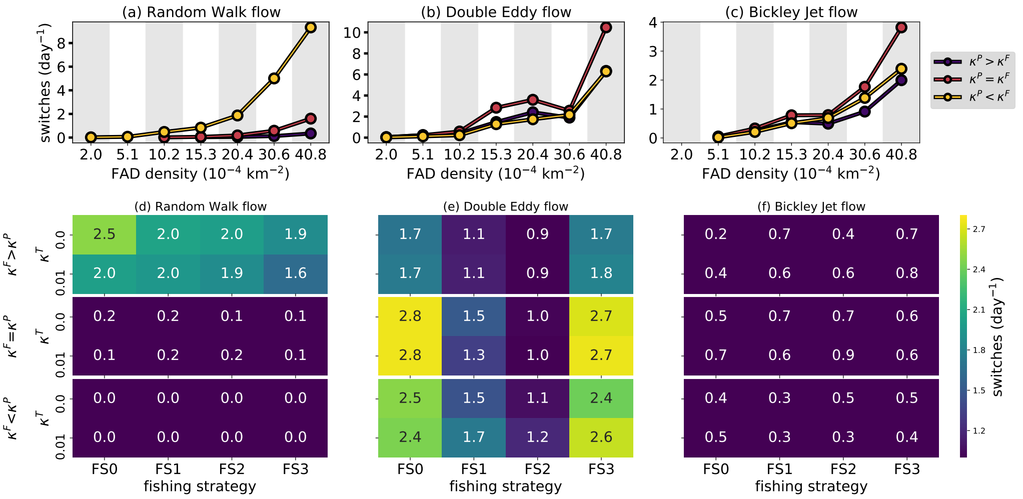


Figure S4: Average frequency of tuna switching from one FAD to another FAD in the Random Walk (left) Double Eddy (middle) and Bickley Jet (right) configuration, averaged over parameter values with FAD ($\kappa^F > \kappa^P$), prey ($\kappa^F < \kappa^P$) and without ($\kappa^F = \kappa^P$) dominant behaviour. (a)-(c) FAD switching at different FAD densities (fishing strategy FS2 and $\kappa^T$=0). (d)-(f) FAD switching for different fishing strategies and values of $\kappa^T$ (FAD density $F$=20).

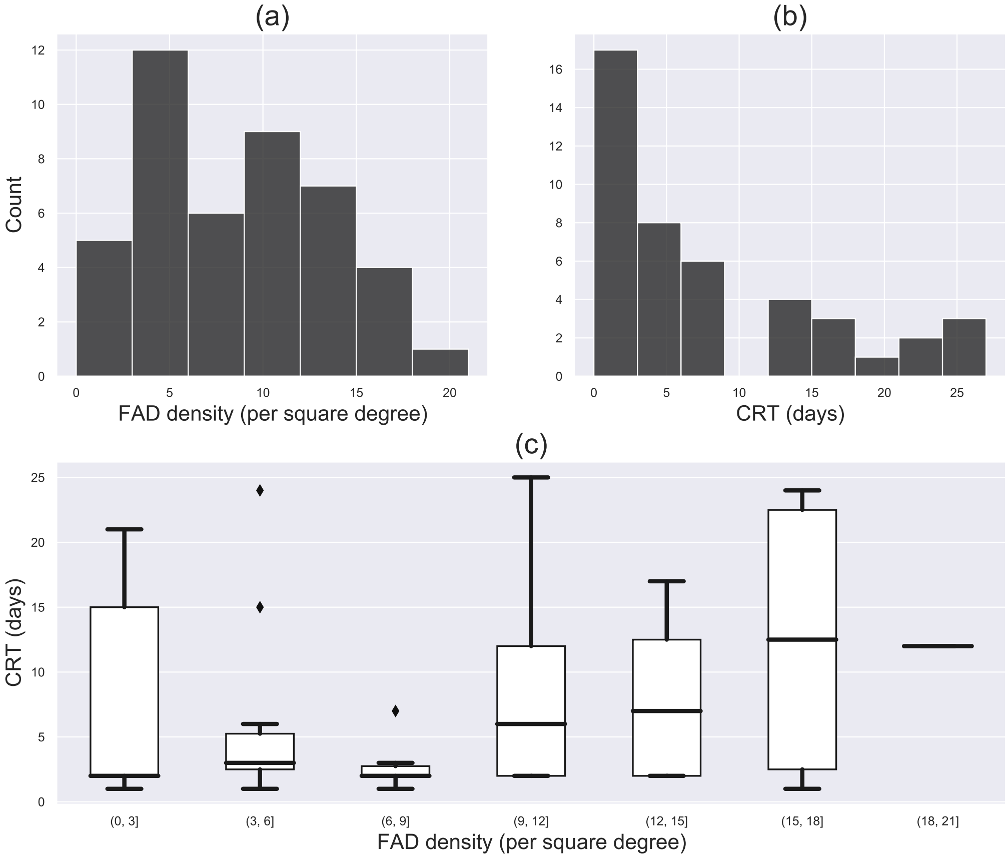


Figure S5: Continuous Residence Times (CRTs) of tuna at FADs, as measured in [9]. Histograms of the number of (a) FAD densities and (b) CRT occurrences in the dataset. (c) Boxplot of CRTs for the same FAD density bins as in (a). Boxplots indicate the CRT median, quartiles, 95% confidence interval and outliers.

### References


[1] R. Olson, C. H. Boggs, Apex Predation by Yellowfin Tuna (Thunnus albacares): Independent estimates from gastric evacuation and stomach contents, bioenergetics, and cesium concentraions, Can. J. Fish. Aquat. Sci. 43 (1986) 1760–1775.

[2] J. J. Magnuson, J. J. Magnuson, Digestion and Food Consumption by Skipjack Tuna (Katsuwonus pelamis) Digestion and Food Consumption by Skipjack Tuna ( Katsuwonus pelamis ), Trans. Am. Fish. Soc. 98 (3) (1969) 37–41.

[3] R. W. Brill, Selective advantages conferred by the high performance physiology of tunas, billfishes, and dolphin fish, Comp. Biochem. Physiol. - A Physiol. 113 (1) (1996) 3–15. doi:10.1016/0300-9629(95)02064-0.

[4] H. S. H. Yuen, Transactions of the American Fisheries Society Swimming Speeds of Yellowfin and Skipjack Tuna, Trans. Am. Fish. Soc. 95 (2) (2011) 203–209. doi:10.1577/1548-8659(1966)95.

[5] J. Scutt Phillips, A. Sen Gupta, I. Senina, E. van Sebille, M. Lange, P. Lehodey, J. Hampton, S. Nicol, An individual-based model of skipjack tuna (Katsuwonus pelamis) movement in the tropical Pacific ocean, Prog. Oceanogr. 164 (February) (2018) 63–74. doi:10.1016/j.pocean.2018.04.007.

[6] G. Moreno, L. Dagorn, G. Sancho, D. Itano, Fish behaviour from fishers' knowledge: The case study of tropical tuna around drifting fish aggregating devices (DFADs), Can. J. Fish. Aquat. Sci. 64 (11) (2007) 1517–1528. doi:10.1139/F07-113.

[7] C. Girard, S. Benhamou, L. Dagorn, FAD: Fish Aggregating Device or Fish Attracting Device? A new analysis of yellowfin tuna movements around floating objects, Anim. Behav. 67 (2) (2004) 319–326. doi:10.1016/j.anbehav.2003.07.007.

[8] Y. Tyutyunov, L. Titova, R. Arditi, Predator interference emerging from trophotaxis in predator-prey systems: An individual-based approach, Ecol. Complex. 5 (1) (2008) 48–58. doi:10.1016/j.ecocom.2007.09.001.

[9] J. Scutt Phillips, B. Leroy, T. Peatman, L. Escalle, N. Smith, Electronic tagging for the mitigation of bigeye and yellowfin tuna juveniles by purse seine fisheries, Tech. Rep. August, Western and Central Pacific Fisheries Commission (2019).
