## Supplementary figures and images for "Modelling of tuna around fish aggregating devices: the importance of ocean flow and prey"

### movie1.gif

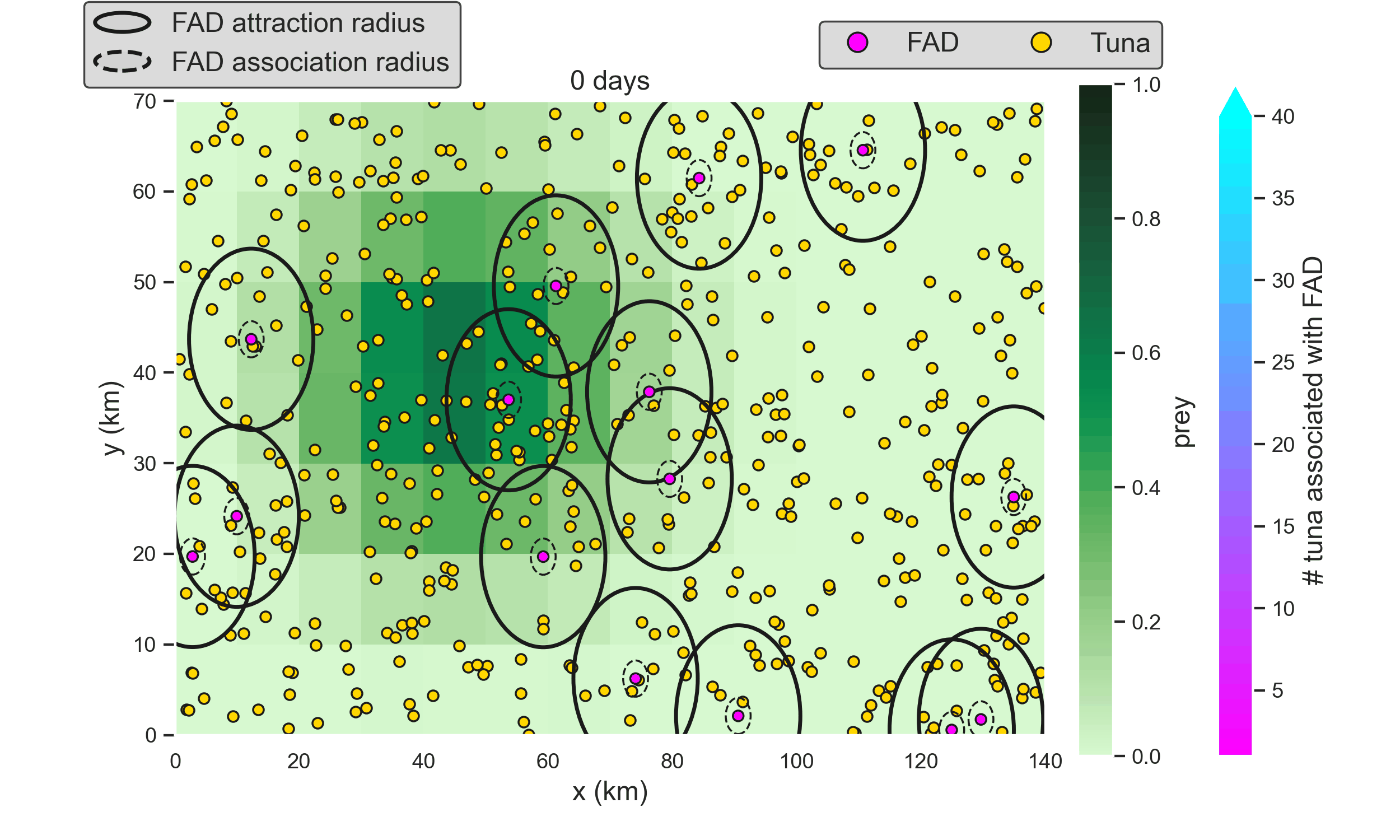

### movie2.gif

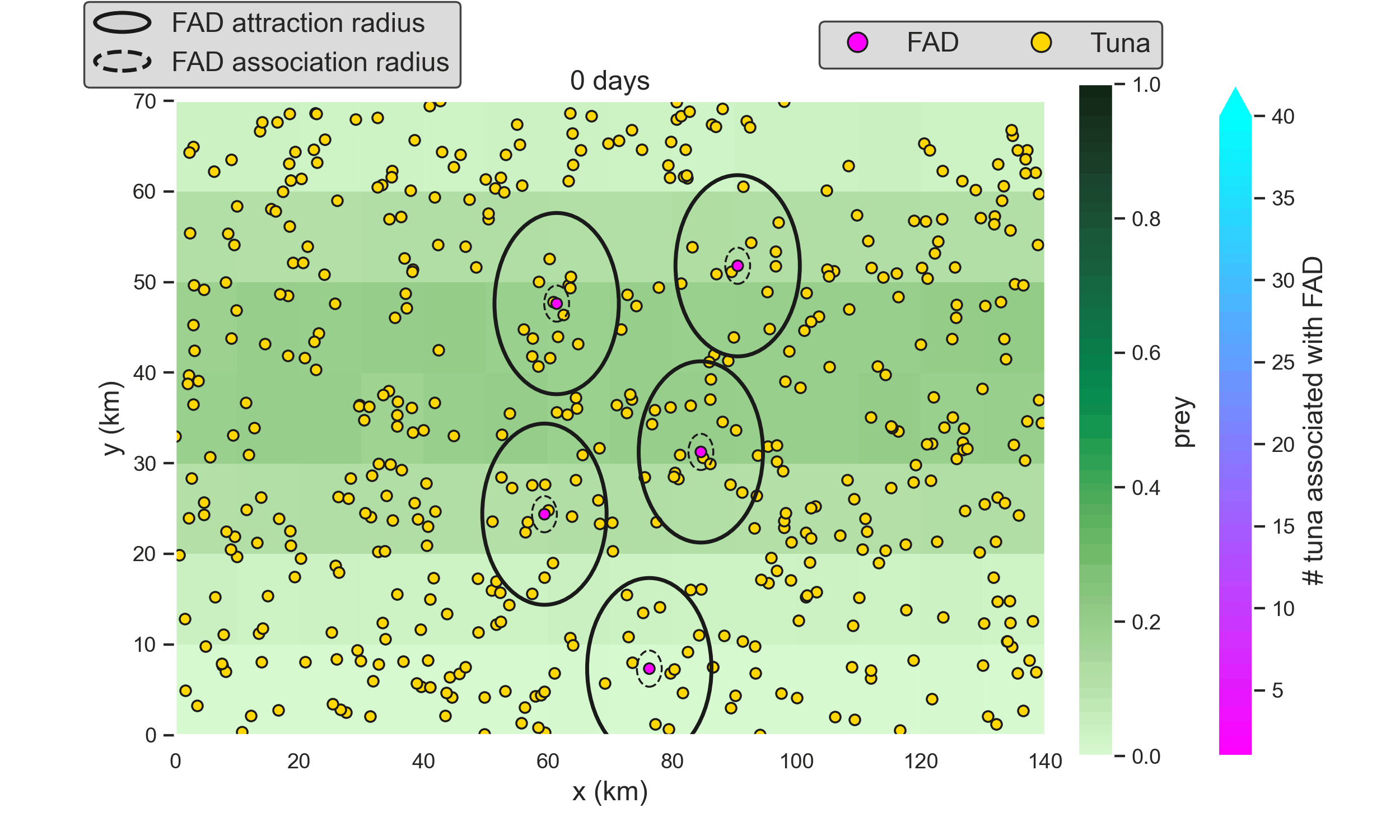

### movie3.gif

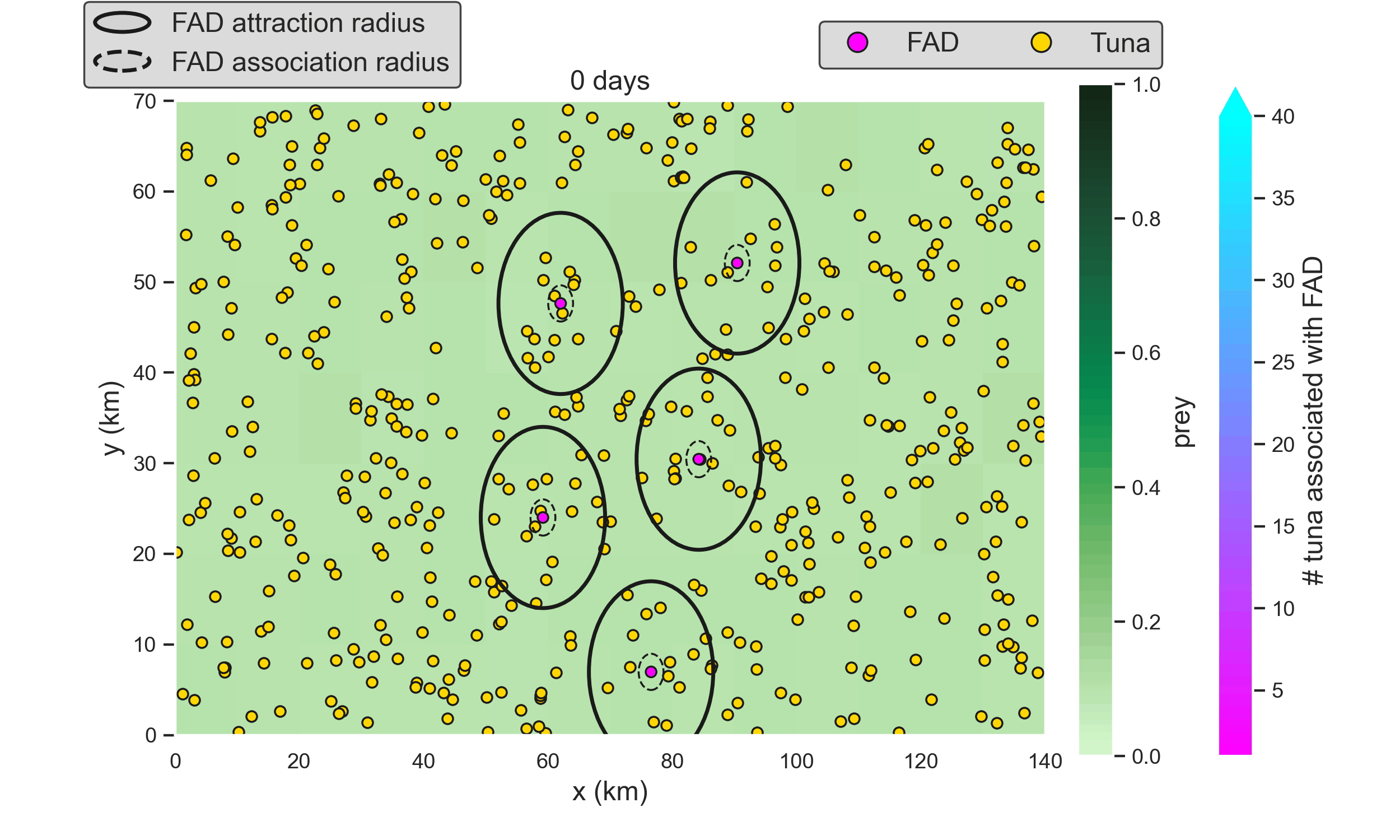

### movie4.gif

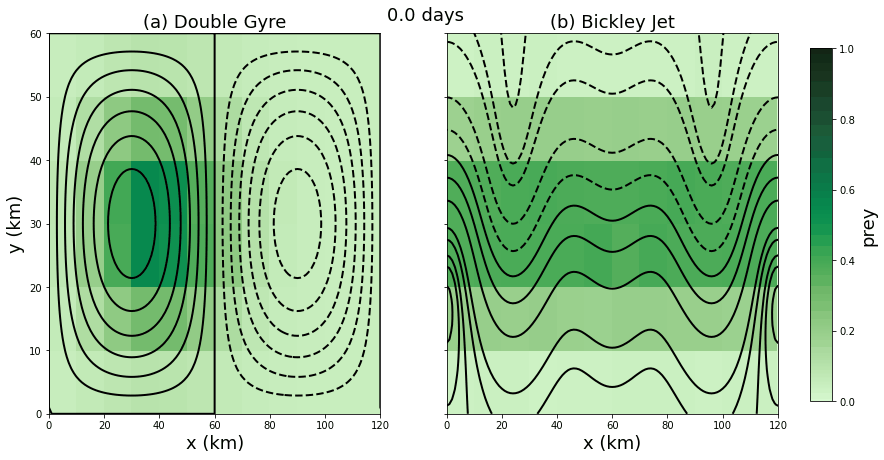
